# A CTGF-YAP regulatory pathway is essential for angiogenesis and barriergenesis in the retina

**DOI:** 10.1101/2020.03.16.994293

**Authors:** Sohyun Moon, Sangmi Lee, JoyAnn Caesar, Sarah Pruchenko, Andew Leask, James A. Knowles, Jose Sinon, Brahim Chaqour

**Author notes:** Author for Correspondence: Brahim Chaqour, B. Chaqour, State University of New York (SUNY) Downstate Medical Center, Dept of Cell Biology, 450, Clarkson Avenue, MSC 5, Brooklyn, NY 11203, USA.

## Abstract

Connective tissue growth factor (CTGF) or CCN2 is a matricellular protein essential for normal embryonic development and tissue repair. CTGF exhibits cell- and context-dependent activities, but the CTGF function in vascular development and permeability barrier is not known. Here we show that endothelial cells (ECs) are one of the major cellular sources of CTGF in the developing and adult retinal vasculature. Mice lacking CTGF expression either globally or specifically in ECs exhibit impaired vascular cell growth and morphogenesis, and blood barrier breakdown. The global molecular signature of CTGF includes cytoskeletal and extracellular matrix protein, growth factor, and transcriptional co-regulator genes such as yes-associated protein (YAP). YAP, itself a transcriptional activator of the CTGF gene, mediates several CTGF-controlled angiogenic and barriergenic transcriptional programs. Re-expression of YAP rescues, at least partially, angiogenesis and barriergenesis in CTGF mutant mouse retinas. Thus, the CTGF-YAP angiomodulatory pathway is critical for vascular development and barrier function.

## INTRODUCTION

Functional vascular networks form as a result of a well-coordinated series of angiogenic events regulating endothelial cell (EC) proliferation, differentiation, polarization, and gene programming to produce stable tubular structures with specific barrier properties (Chow and Gu, 2015; Zhao et al., 2015). In the retina, angiogenesis occurs during postnatal stages when ECs from the brain invade the optic nerve, emerge from it and spread over and within the retinal neuroepithelium (Fruttiger, 2007). This angiogenic process is coordinated with the simultaneous formation of a blood retinal barrier wherein the paracellular permeability between adjacent ECs and transcellular transport across ECs are tightly regulated by junctional protein complexes and transporter proteins, respectively (van der Wijk et al., 2019). Cells within the cohesive vascular wall have cadherin-based adhesions at cell–cell junctions and integrin-based focal adhesions to cell-extracellular matrix (ECM) contacts. ECM proteins, in particular, form a network of fibrillary and fibrous proteins and glycans that are critical for all aspects of vascular growth and regeneration (Bishop, 2015). They provide both anchorage points to the cells and chemical cues for directional migration, morphogenesis, and stability. In addition, EC-ECM interactions contribute to the acquisition of EC barrier properties appropriate for the nervous system (Scott et al., 2010; Segarra et al., 2018). However, the diversity of both the matrisome components (which constitute >1% of the proteome) and the mechanisms controlling the synthesis, composition and remodeling of ECM proteins is suggestive of an intricate level of complexity for the control angiogenesis and barriergenesis (Bou-Gharios et al., 2004).

Connective tissue growth factor (CTGF) also named cellular communication network (CCN) 2 is a candidate ECM protein whose precise function in the vascular matrix is largely unknown. CTGF was originally isolated by differential screening of cDNA libraries prepared from HUVECs and NIH 3T3 fibroblasts (Ryseck et al., 1991). The protein was named CTGF because of its mitogenic activity vis-à-vis fibroblasts and cultured ECs. CTGF is also referred to as cellular communication network 2 (CCN2) by virtue of its multimodular and structural analogy to the CCN family of proteins (Krupska et al., 2015; Perbal et al., 2018). The primary CTGF translational product is a ∼40 kDa protein that contains 38 conserved cysteine residues dispersed throughout four distinct structural modules. CTGF elicits its biological activities through binding to various cell surface receptors including integrin receptors, cell surface heparan sulphate proteoglycans, low-density lipoprotein receptor-related proteins, and TrkA in a cell type- and context-dependent manner (Gao and Brigstock, 2004; Lau, 2016). It was suggested that such interactions enable CTGF to regulate a variety of cellular functions including cell adhesion, proliferation, migration, differentiation, survival, and ECM synthesis. In addition, *in vitro* assays showed that the CTGF N-terminal and C-terminal moieties interact with glycoproteins, proteoglycans, growth factors and proteases although the *in vivo* significance of such interactions is not well understood (Dean et al., 2007; Hashimoto et al., 2002; Inoki et al., 2002; Pi et al., 2011). Global CTGF deficiency in mice demonstrated the importance of CTGF in cardiovascular and skeletal development, as CTGF-null mice exhibited defects in basic lung development and failed thoracic expansion, leading to perinatal lethality (Ivkovic et al., 2003). Although no obvious vascular alterations were observed in CTGF-deficient mice during the initial formation of the primitive blood vessels, deficiencies in the endocrine cell lineage and reduced growth plate angiogenesis were noted. In adults, CTGF plays an important role in wound repair and CTGF protein levels correlate with many vascular and inflammatory diseases such as arthritis, diabetic nephropathy and retinopathy (Chintala et al., 2012; Leask et al., 2002; Nguyen et al., 2008; Praidou et al., 2010; Tang et al., 2018). In a rat model of glomerulonephritis, CTGF levels were elevated in areas of crescentic extracapillary proliferation, periglomerular fibrosis, and in interstitial foci (Gupta et al., 2000; Toda et al., 2017). While these *in vivo* studies suggested a potentially important role of CTGF in physiological and pathological angiogenesis, the specific function and mechanisms whereby CTGF regulates blood vessel development and function remain to be investigated.

Here we provide the first evidence that CTGF directly regulates retinal tissue vascularization and blood barrier integrity. Our data indicate that loss of CTGF function impairs a genetic angiogenic program involved in vessel sprout morphogenesis and branching and barrier integrity. We further provide data about exemplar CTGF target genes such as YAP, which when re-expressed in the vasculature, rescues, at least in part, transcriptional programs associated with CTGF deficiency. Our results shed light on the genetic landscape responsible for CTGF-dependent regulation of proper vessel formation and function.

## RESULTS

### Robust CTGF Expression in Developing and Adult Vasculature

The retina is a complex neurovascular tissue organized into three cellular and two synaptic layers supported by a tripartite intraretinal vascular network. In mice, the retina is supported by three interconnected vascular layers that develop postnatally, making this model particularly useful for assessing the role of genes potentially relevant to human vascular development and function (Lee et al., 2019; Lee et al., 2017a; Lee et al., 2017b). We first examined CTGF expression in the retina of newborn and adult mice. CTGF transcript levels increased progressively between P0 and P24 when the capillary plexuses were forming and plateaued thereafter (Fig. 1A). CTGF protein levels similarly increased as retinal vascular plexuses invade the retina and expression of the intact protein persists in the adult vasculature (Fig. 1B-C). To determine the precise cellular sources of CTGF, we used a CTGF-GFP reporter transgenic mouse line in which the GFP reporter gene was placed downstream of a large CTGF promoter segment (>100-kb). As shown in Fig. 1D, the reporter expression was found mainly in the expanding vascular network at P2 and P6 and persisted throughout the postnatal and adult periods. Further detailed analysis revealed a robust expression of CTGF in ECs of the sprouting primary capillary plexus. Endothelial tip cells with their filopodial extensions expressed little or no CTGF-GFP signal (Fig. 1E), whereas the trailing stalk ECs showed a strong CTGF-GFP signal indicating that CTGF potentially regulates stalk cell function including proliferation, lumenization, and stabilization. In addition, as the vascular tree developed, the CTGF-GFP signal spread widely into in NG2-positive mural cells (i.e., pericytes) of small and larger vessels (Fig. 1F). EC and pericyte expression of CTGF persisted in adult retinal tissue as well. The brain vasculature, which is similar in its complexity and properties to the retinal vasculature, exhibited a strong expression of the CTGF-GFP marker as well (Fig. S1A). In the developing retina, detailed analysis or tissue cross-sections revealed CTGF-GFP signal in glutamine synthase (GS)^+^ Muller cells and Iba-1^+^ microglia particularly those of the inner layer (Fig. 1G-H). Expression of CTGF was not associated with astrocytes (not shown) even though these cells are a major source of vascular endothelial growth factor (VEGF), a *bona fide* regulator of CTGF expression in cultured cells (Asano et al., 2019; Chappell et al., 2019; Nakamura-Ishizu et al., 2012; Suzuma et al., 2000). Thus, CTGF exhibits heterogeneous and spatiotemporal patterns during vessel morphogenesis and a sustained expression in the adult vasculature of the central nervous system.

**Fig 1.**
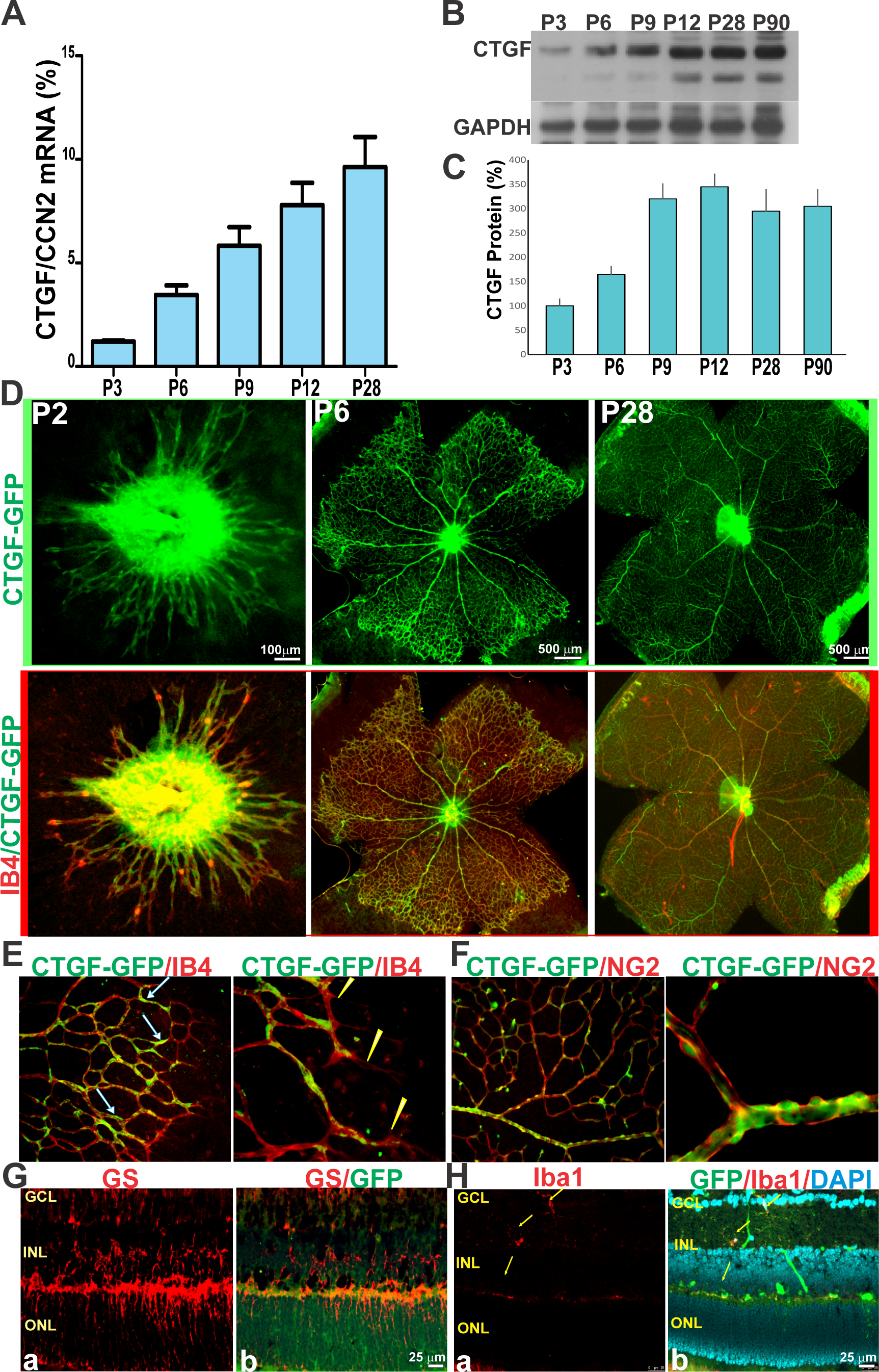
Expression pattern of CTGF in the developing and adult blood vessels in mouse retina. (A-C) Analysis of CTGF expression at the RNA (A) and protein (B) levels by real time PCR and Western blotting, respectively, in retinal lysates during postnatal development of the retinal vasculature. CTGF mRNA levels were normalized to those of GAPDH. CTGF protein bands were quantified by densitometric scanning (C). (D) Unstained (upper panels) and isolectin B4 (IB4; red) stained flat-mounted retinas of P2, P6 and P28 CTGF-GFP reporter mice. Merged images in the lower panels show vascular localization of the CTGF-GFP signal. (E-F) High magnification images of vascular front of flat-mounted retinas of CTGF-GFP reporter mice stained with IB4 and NG2 respectively. Arrows and arrowheads in (E) indicate CTGF-GFP reporter signals in endothelial stalk cells and lack thereof in tip cells, respectively. Note the expression of the CTGF-GFP signal in NG2^+^ pericytes in (F). (G-H) Transverse retinal section of CTGF-GFP mice stained with glutamine synthase (GS) and Iba1. GS or Iba1 staining alone or merged with GFP are shown. Retinal layers indicated are ganglion cell layer (GCL), inner nuclear layer (INL), and outer nuclear layer (ONL). Arrows indicate CTGF-GFP-Iba-1-positive cells.

### Loss of CTGF Expression Alters Vessel Morphogenesis

As a secreted protein, CTGF exerts autocrine and paracrine actions because it localizes not only within the interstitial ECM but also pericellularly, due to its strong heparin-binding activity (Chaqour, 2013; Kireeva et al., 1997). To gain new insight into the function of CTGF produced by vascular cells, we examined the vascular phenotypes associated with either global, EC- or pericyte-specific loss of CTGF function. We used a CTGF mutant strain in which loxP sites had been engineered to flank exons 1 and 2 (Liu et al., 2011) (Fig. S2A). CTGF^flox/flox^ mice were crossed with transgenic Cre mice bearing an inducible Cre recombinase under the control of either the ubiquitin C (UBC), Cdh5, or Cspg4 promoter to produce mouse mutants with global, EC-, and pericyte-specific deletion of CTGF (hereafter referred to as UBCΔCTGF, Cdh5ΔCTGF and CspgΔCTGF), respectively (Fig. S2B). The expression of CTGF was quantified by quantitative PCR to confirm its knockdown after three consecutive 4-hydroxy tamoxifen (4HT) injections at P1, P2 and P3. WT and mutant mouse retinal vascular phenotypes were examined at P7 when the vasculature is still growing but it had also acquired tissue- and barrier-specific properties to support organ function. UBC-, Cdh5- and Cspg4-CreER^T2^-recomination with floxed alleles effectively reduced CTGF mRNA levels by >85%, 55% and 50% compared to wild-type (WT) (i.e., CTGF^flox/flox^) mice respectively (Fig. S2C). CTGF protein levels were consistent with those of CTGF mRNA in WT and mutant mice (Fig. S2D). The less complete recombination of the Cdh5- and Cspg4-CreER^T2^ alleles is likely due to the expression of CTGF by cells other than ECs and pericytes, respectively. Retinas from 4HT- or corn oil-injected littermate UBC-CreER^T2^, Cdh5(PAC)-CreER^T2^ and Cspg4-CreER^T2^ exhibited a vascular phenotype identical to CTGF^flox/flox^ mouse retinas and were used as controls in all experiments. As shown in Fig. 2A-D, global loss of CTGF function resulted in significant reduction of microvessel density, vascular branching and sprouting vessels at the vascular front. While arteriovenous differentiation and vascular expansion to the retinal edge were not affected, the vascular network was partially shaped into rudimentary arterioles, venules, and capillaries. BrdU^+^ cell count at the angiogenic front showed a 45% reduction of cell proliferation (Fig. 2E). Pericyte coverage (normalized to vascular area) was seemingly unaffected in UBCΔCTGF mutant mouse vessels suggesting that CTGF loss had no apparent effect on vessel coverage by mural cells (Fig. 2F). Similarly, EC-specific CTGF deletion induced a significant decrease in vascular area, branching points, and cell proliferation, and simultaneous increase of vascular lacunarity although these alterations were slightly milder than those of the UBCΔCTGF mutants (Fig. 2G-K). Conversely, the CspgΔCTGF retinal vasculature closely resembled the WT control with respect to vascular morphology, architecture and density (Fig. 2L-P), indicating that pericyte-derived CTGF has no effect on retinal vascular development. Thus, CTGF deficiency in the endothelium recapitulates, at least in part, the global loss-of-function phenotype.

**Fig 2.**
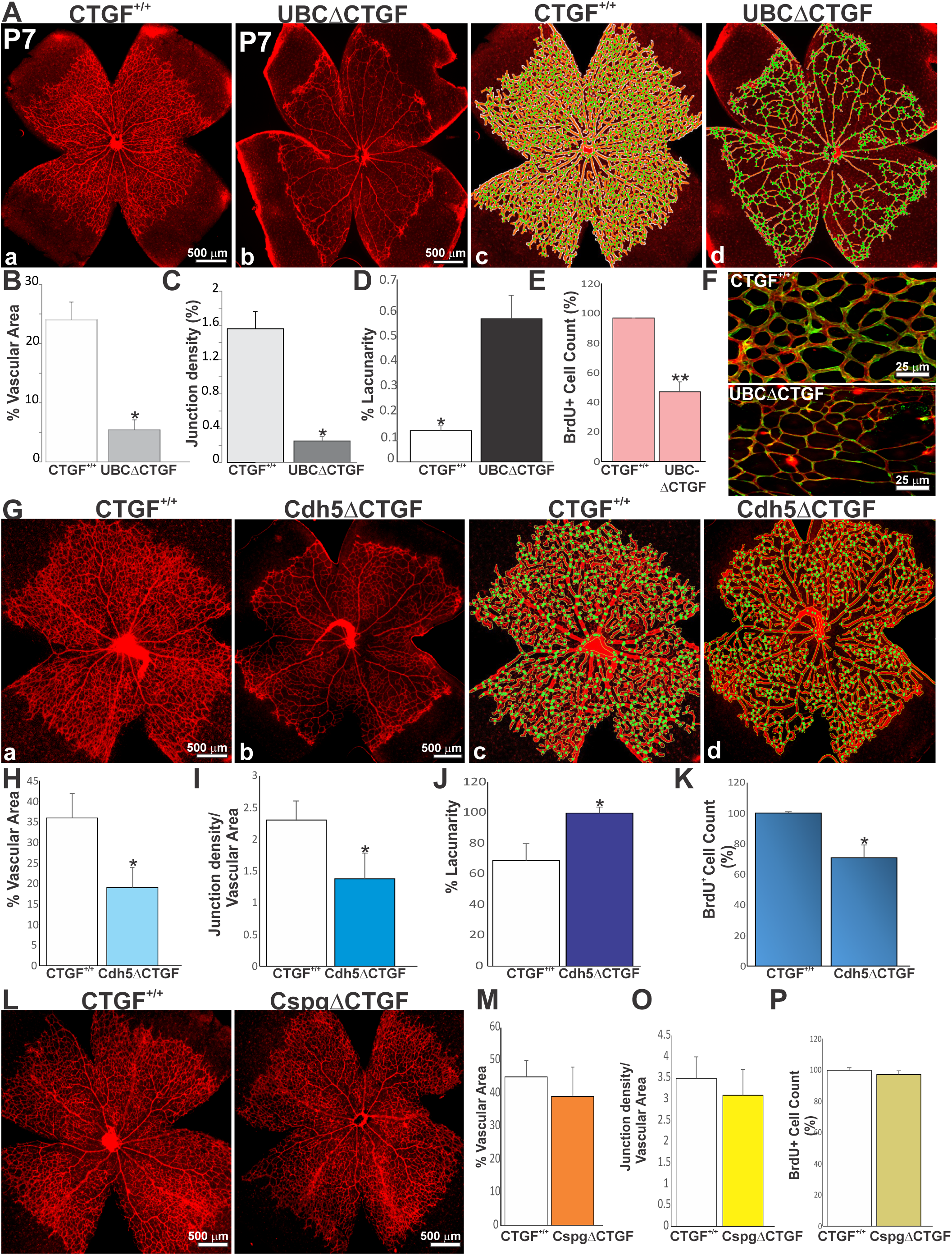
Loss of CTGF function altered normal vessel growth and development. (**A**) IB4-stained whole retinal mounts of WT (a) and UBCΔCTGF (b) mice. Quantitative analyses of vascular parameters were performed with AngioTool software. The vasculature outline, skeleton, and branching points were denoted in white, red, and green, respectively (c-d). (**B-D**) Graphical representations of the changes in vascular surface, junction density and lacunarity in WT and UBCΔCTGF mouse retinas. *, p <0.001 *vs* CTGF^+/+^ (n=5). (**E**) Number of BrdU^+^ proliferating ECs per retinal area unit. **, p <0.05 *vs* CTGF^+/+^ (n=5). **(F)** Representative immunofluorescence images of dual IB4 (red) and NG2 (green) staining of whole mount retinas of CTGF^+/+^and UBCΔCTGF mice. (**G**) IB4-stained whole retinal mounts of wild-type CTGF^+/+^ (a) and Cdh5ΔCTGF (b) mice and associated AngioTool analysis [shown in (c) and (d)]. (**H-K**) Graphical representations of the changes in vascular surface, junction density and lacunarity and BrdU^+^ cell count. *, p <0.05 *vs* CTGF^+/+^ (n=5). **(L-P)** Effects of pericyte-specific deletion of CTGF on the retinal vascular phenotype. Vascular parameters and cell counts were determined as described in (A-E).

### Loss of Barrier Function Following Global and EC-Specific Deletion of CTGF

During vessel development, ECs tightly coordinate angiogenesis with barrier function formation. The latter involves the expression and engagement of tight junction (TJ) and transporter proteins that limit paracellular and transcellular permeability between and across ECs, respectively (Dejana et al., 2008). To assess the effects of CTGF on barrier function, we examined extravasation of retro-orbitally-injected fluorescein isothiocyanate (FITC)-albumin, which does not traverse the vascular barrier under physiological conditions. In WT mice, the injected tracer was confined to the intravascular space of the retina (Fig. 3A). However, both UBCΔCTGF and Cdh5ΔCTGF mutant mice exhibited extensive vascular leakage, indicated by a diffuse hyperfluorescent background and patchy hyperfluorescence in the extravascular space. Conversely, pericyte-specific deletion of CTGF did not produce overt signs of FITC-albumin extravasation, which, in most cases, was similar to that of a WT mouse retina. Similarly, EB leakage from retinas with either global or EC-specific deletion of CTGF was 4- to 9-fold higher than that of wild-type controls. There was no significant difference between WT and CspgΔCTGF mice with respect to EB leakage, which is consistent with the FITC-albumin results (Fig. 3B). To determine whether the expression of CTGF is required for barrier maintenance as well, we examined retinal vascular permeability in adult (P36) mice following 4HT injection a week prior. Cdh5ΔCTGF retinas showed numerous bright extravascular spots due to FITC-albumin leakage from the retinal vasculature (Fig. 3C). Likewise, FITC-albumin extravasation is observed in the brain vasculature of the mutant mice indicating a defective blood brain barrier (Fig. S1B). In contrast, age-matched littermates showed no tracer leakage in the retinal and brain parenchyma. Vascular leakage index was 41% higher in CTGF mutant compared to age-matched WT retinas (Fig. 3D), confirming the importance of CTGF signals in blood retinal barrier maintenance.

**Fig 3.**
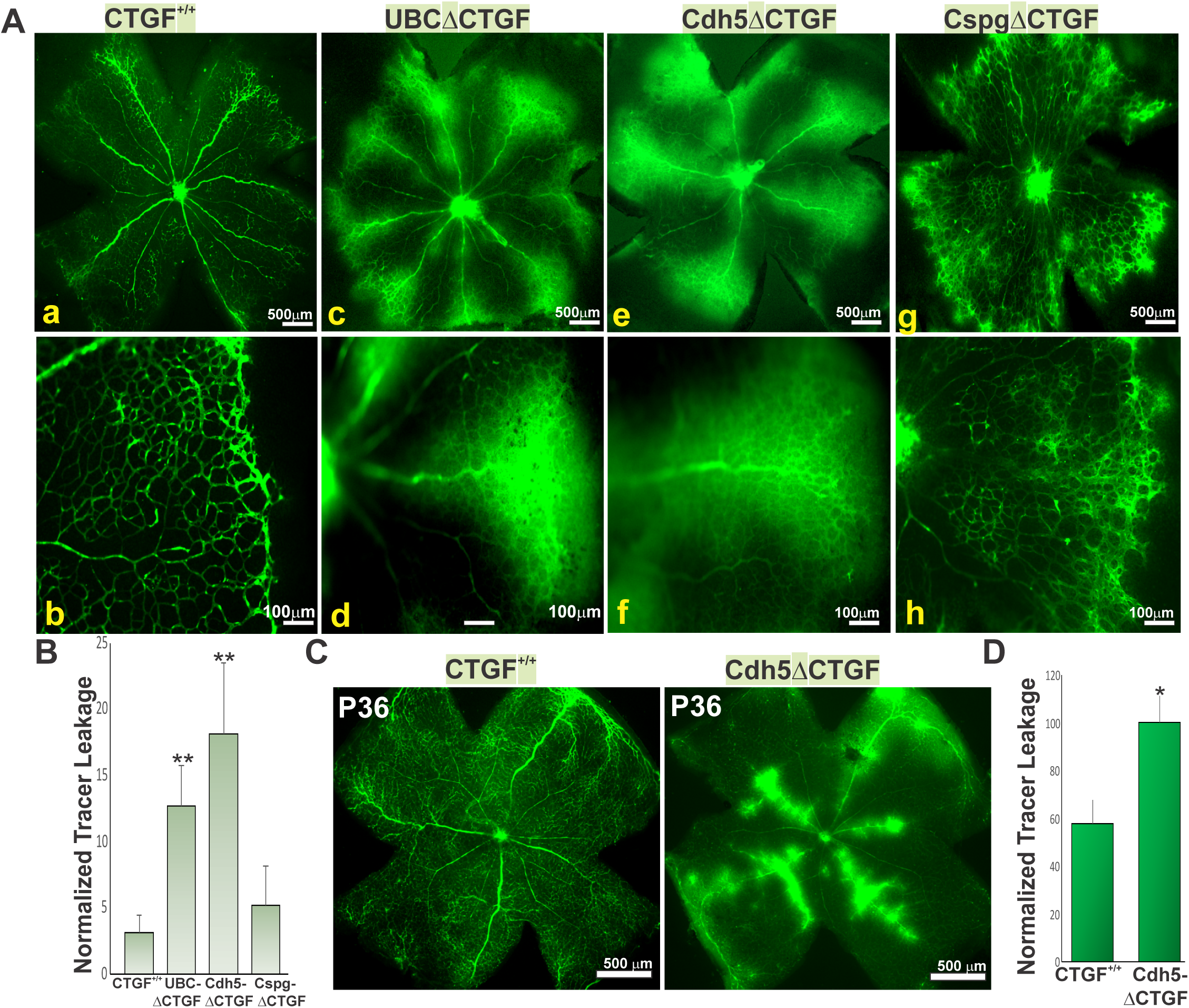
CTGF deficiency induced vascular leakage in the retina. (A) FITC-albumin-injected retinal flat mounts from CTGF^+/+^ (a, b), UBCΔCTGF (c, d), Cdh5ΔCTGF (e, f) and CspgΔCTGF (g, h) mice. Note the absence of FITC-albumin extravasation in CTGF^+/+^ and CspgΔCTGF and hyper-fluorescence of FITC-albumin in the retinal parenchyma of UBCΔCTGF and Cdh5ΔCTGF mice. (B) Vascular permeability index was normalized to total vascular area. (C) FITC-albumin-injected retinal flat mounts from CTGF^+/+^ and Cdh5ΔCTGF adult (P36) mice that have received three consecutive injection of 4HT a week prior. (D) Vascular permeability index measured as described in B. Values are means ± SEM (n=3). Values in Cdh5ΔCTGF mice were set to 100% to facilitate comparisons among animals.

### Molecular Signature of CTGF During Retinal Vascular Development

The formation of an organized retinal vasculature depends on signaling among different vascular and non-vascular cellular components of the retina including ECs, mural cells, neurons, glia, and immune cells. As an ECM protein, CTGF signals may affect the interactions among all these cellular components. To determine the global genetic bases of CTGF-dependent regulation of vascular growth and barrier function, we examined the transcriptomic differences between CTGF^+/+^ and UBCΔCTGF mutant mouse retinas through RNA-Seq. Following three consecutive 4HT administration from P1 onwards, retinas from three different litters of CTGF^flox/flox^ and CTGF^flox/flox^-UBC-CreER^T2^ intercrosses were harvested at P7 and processed for total RNA extraction and gene profiling. Unsupervised hierarchical clustering and principal component analyses (PCA) demonstrated clear segregation and reproducibility of the obtained gene expression profiles for WT and UBCΔCTGF mutants (Pearson’s correlation coefficient ρ = 0.94) (Fig. S3A-C). The candidate CTGF-regulated genes encode proteins with a wide range of biological activities, likely reflecting the large diversity of cell types and circuits in the retina (Fig. S3D). Genes that encode proteins for the biosynthesis of RNA, macromolecules, nucleobase-containing compounds, and neurogenesis and RNA polymerase transcription were highly differentially expressed between WT and mutant retinas.

To expand our RNA-Seq data analysis beyond the initial cut-off parameters, which often are subjective, arbitrary and not biologically justified, we used gene set enrichment analysis (GSEA) to map all detected unfiltered genes to defined gene sets (e.g., pathways), irrespective of their individual change in expression (Subramanian et al., 2005). We identified 209 significantly changed genes (133 downregulated and 76 upregulated genes) in UBCΔCTGF compared with CTGF^+/+^ (absolute log_2_ fold change >0.5, p < 0.05) (Fig. 4A). When the normalized data of the CTGF^+/+^ and UBCΔCTGF transcriptomes were compared to 1, 454 GO gene sets in the GSEA Molecular Signatures Database (MSigDB)(Liberzon et al., 2015), twenty gene sets were significantly enriched in either the CTGF^+/+^ or UBCΔCTGF group. Of these, thirteen were closely related to neuronal progenitor differentiation linked to CRX expression, cell cycle regulation (i.e., cyclins, p53, SRC, Kras, YAP), cell survival (i.e., Akt), and inflammation pathways (i.e., IL-15, NF-kB) (Fig. 4B). Gene sets of the YAP, PDGF, and mTOR pathways were similarly downregulated in UBCΔCTGF. Other core-enriched genes are involved in the regulation of tip cell differentiation, angiogenesis, ECM, and ECM metabolic pathways (Fig. 4C-E). In addition, CTGF deficiency resulted in downregulation of angiopoietin 1, ROBO1, integrin β1, Slit-1, Ephrin B2, Ephrin Receptor B4, and Dll-1, and upregulation of integrin β1 binding protein 1 and thrombospondin 1. Downregulated ECM protein genes include laminin 4, elastin, SPARC, fibrillin-2, glypican 5, collagen 4A1, and tenascin-C, whereas vitronectin, Col17A1, Col16A1 and Col5A3 genes were upregulated upon CTGF deletion. Genes encoding growth factors such as IGF-1, pleiotrophin, opticin, LTBP-1, TGF-β1, TGF-β2 and FGF-10 were downregulated in UBCΔCTGF mutant mice. These results are suggestive of an important role of CTGF in regulating angiogenesis-related processes such as cell adhesion, migration and guidance, as well as cell-matrix interaction. Importantly, the expression of several transcriptional regulator genes (e.g., YAP, TEAD, NFAT5, SRF) was dysregulated by loss of CTGF function. Transcription factors are endpoints of a network of signaling pathways enabling the cells to process signals they receive simultaneously from many different receptors. The multitude of transcription factors affected by CTGF deletion suggests that CTGF signals impact global mechanisms of gene regulation.

**Fig. 4.**
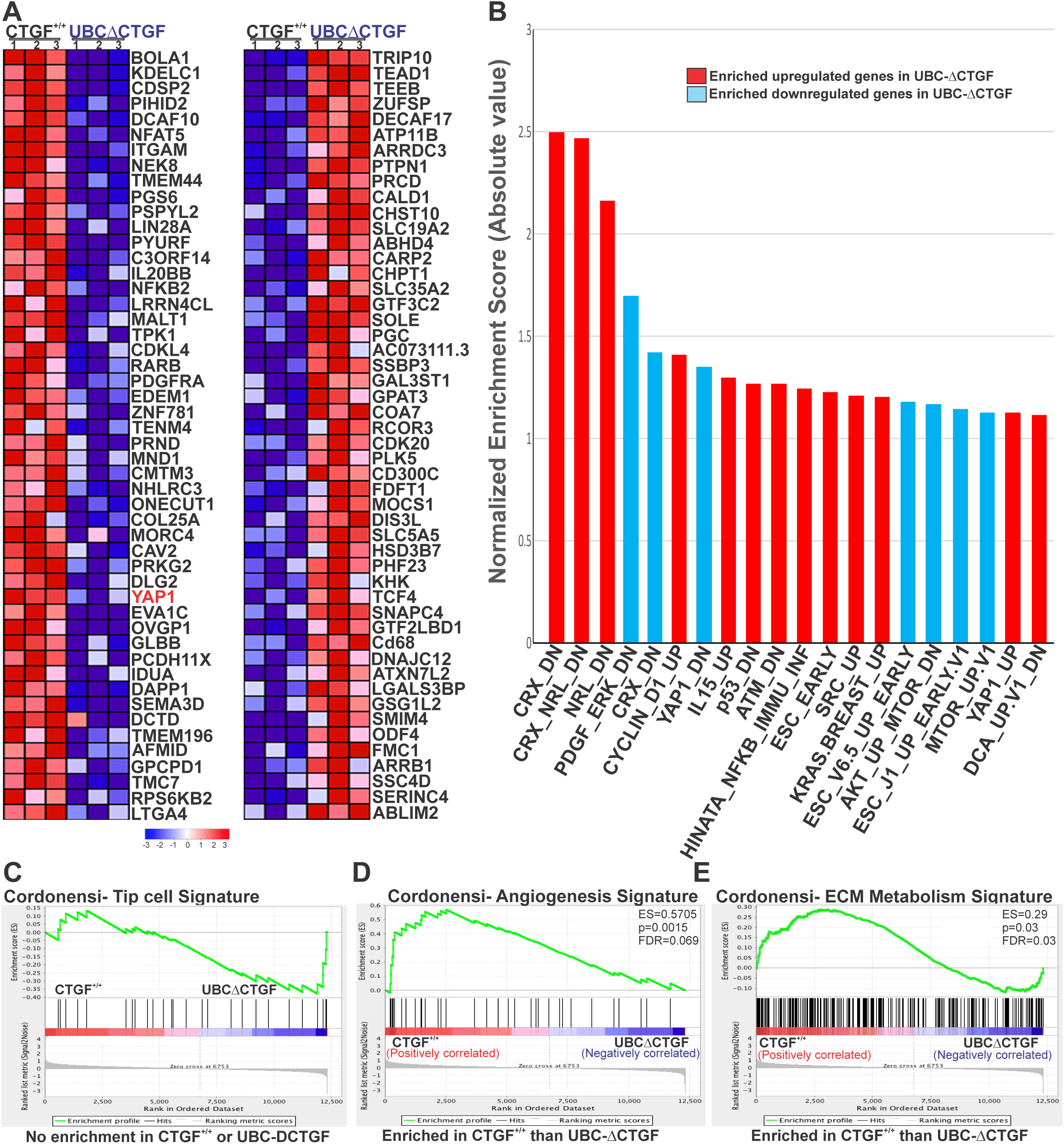
Identification of gene sets enriched in CTGF^+/+^ and UBCΔCTGF phenotype as determined by GSEA. (A) GSEA-generated heatmaps of defined gene sets enriched at the top of a gene list ordered on the basis of expression differences between CTGF^+/+^ and UBCΔCTGF mouse retinas. Range of colors (red to blue) indicates the range of expression values (high to low). (B) Pathway enrichment analysis of differentially expressed genes in UBCΔCTGF mouse retina. The clusters with functional terms that reached significance of p ≤ 0.05 are shown. Online functional databases were used to extract each term. (C-E) Enrichment plot of the gene set of tip cell, angiogenesis and ECM/cell adhesion pathways. Genes that appear before or at the peak are defined as core enrichment genes for that gene set. Genes whose expression levels are most closely associated with the CTGF^+/+^ group are located at the left while genes from the UBCΔCTGF gene set within the ranked list are located at the right.

### CTGF-Dependent Regulation of YAP Expression Affects Blood Vessel Growth and Morphogenesis

YAP was among the most remarkable transcriptional regulators and key angiogenic factors that was differentially expressed as a result of CTGF deletion (Fig. 4A and Fig. S2D). Like CTGF, YAP expression has been reported to increase in the retinal vasculature at an early stage of development (Choi et al., 2015; Neto et al., 2018). YAP is also required for the VEGF-VEGFR2 signaling axis to promote angiogenesis (Wang et al., 2017). VEGF-induced YAP activation leads to the activation of Src family kinases and successive cytoskeletal remodeling, which orchestrates cellular decisions of proliferation, differentiation, cell shape, and polarity (Kim et al., 2017). To investigate the role of YAP in the underlying mechanisms of CTGF-dependent regulation of angiogenesis, we determined whether the transcriptomic changes associated with CTGF deficiency correlated with those caused by YAP deletion (or overexpression). Using GSEA and previously reported data for YAP overexpression in the Msig (Liberzon et al., 2015), we found that numerous genes that were upregulated upon YAP stimulation were decreased by CTGF deletion (e.g., Filamin A, ASAP1, CRIM1, SLIT2) (Fig. 5A-C). Reversibly, genes that were downregulated upon YAP stimulation were upregulated upon CTGF deletion (e.g., Cox8A, RAB40C). Since CTGF is itself a YAP target gene, the activity of CTGF and YAP are mutually regulated. For CTGF target gene data validation, we analyzed 11 differentially expressed YAP network genes using quantitative real time RT-PCR. The expression of 9 of the 11 genes was fully consistent with their RNA-Seq expression profiles and trends (p<0.05) (Figure 5A), suggesting that the effects of CTGF were, at least in part, mediated through YAP. Thus, the CTGF-dependent transcriptional program is linked to YAP regulatory activities.

**Fig. 5.**
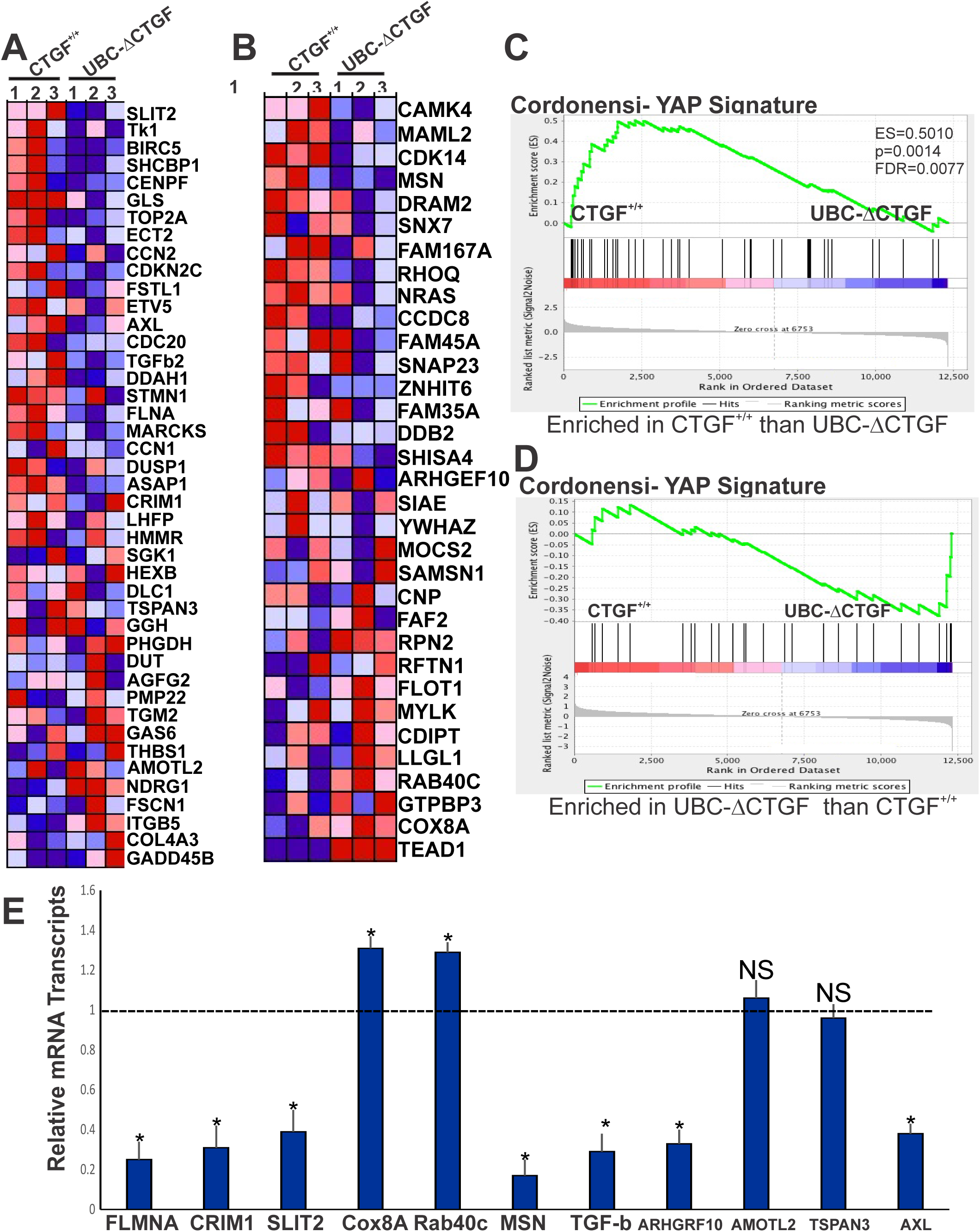
CTGF-dependent transcriptional program targets primarily YAP network genes. (A-B) Heatmaps of the top 32-43 enriched genes identified by GSEA of P7 mouse retina showing differential regulation of YAP signature genes in UBCΔCTGF compared with those in CTGF^+/+^ mice. Enrichment of CTGF signature with genes up-regulated in MCF10A cells (breast cancer) over-expressing YAP gene and YAP conserved signature/Hippo pathway genes are shown in (A) and (B) respectively. (C-D) Running enrichment score for the gene set analyses shown in (A) and (B) as the analysis walks along the ranked list. (E) Relative mRNA levels of YAP target genes in retinal lysates from CTGF^+/+^ and UBCΔCTGF mouse retinas. The mRNA levels in CTGF^+/+^ mice were set to 1. Each measurement was performed in triplicate. (*, p <0.05, *n*=3).

To further assess the extent of YAP involvement in CTGF-dependent effects on vascular development, we determined whether ectopic re-expression of YAP could rescue the vascular sprouting phenotype associated with CTGF deletion. To this end, YAP was ectopically expressed through adeno-associated virus type 6 (AAV6)-mediated gene transfer (Fig. 6A). Retro-orbital injection of a control AAV6-GFP vector into P1 mouse pups produced efficient expression of GFP in ECs of retinal blood vessels at P7 (Fig. 6B). Quantitative analyses of the retinal vascular phenotype at P7 showed that AAV-mediated re-expression of YAP increased vascular area and junction density in UBCΔCTGF compared to those injected with AAV-control gene (i.e., AAV6-luc) (Fig. 6C-E). Similarly, BrdU-positive cell count was significantly increased upon re-expression of YAP in CTGF-deficient mice, which is consistent with increased cell proliferation in sprouting vessels (Fig. 6F-G). Quantitative PCR-based analyses of mRNA contents showed that YAP re-expression primarily affected genes involved in cell adhesion and migration (e.g., CCN1, TSPAN, CRIM1), proliferation (e.g, ECT2, SLIT2), differentiation (e.g., HEY1) and ECM remodeling (e.g., TIMP3) (Fig. 6H). These gene targets are under either direct or indirect regulation by YAP (Kim et al., 2016; Kim et al., 2015b), suggesting that YAP acts as a relay for CTGF-dependent control of EC growth.

**Fig. 6.**
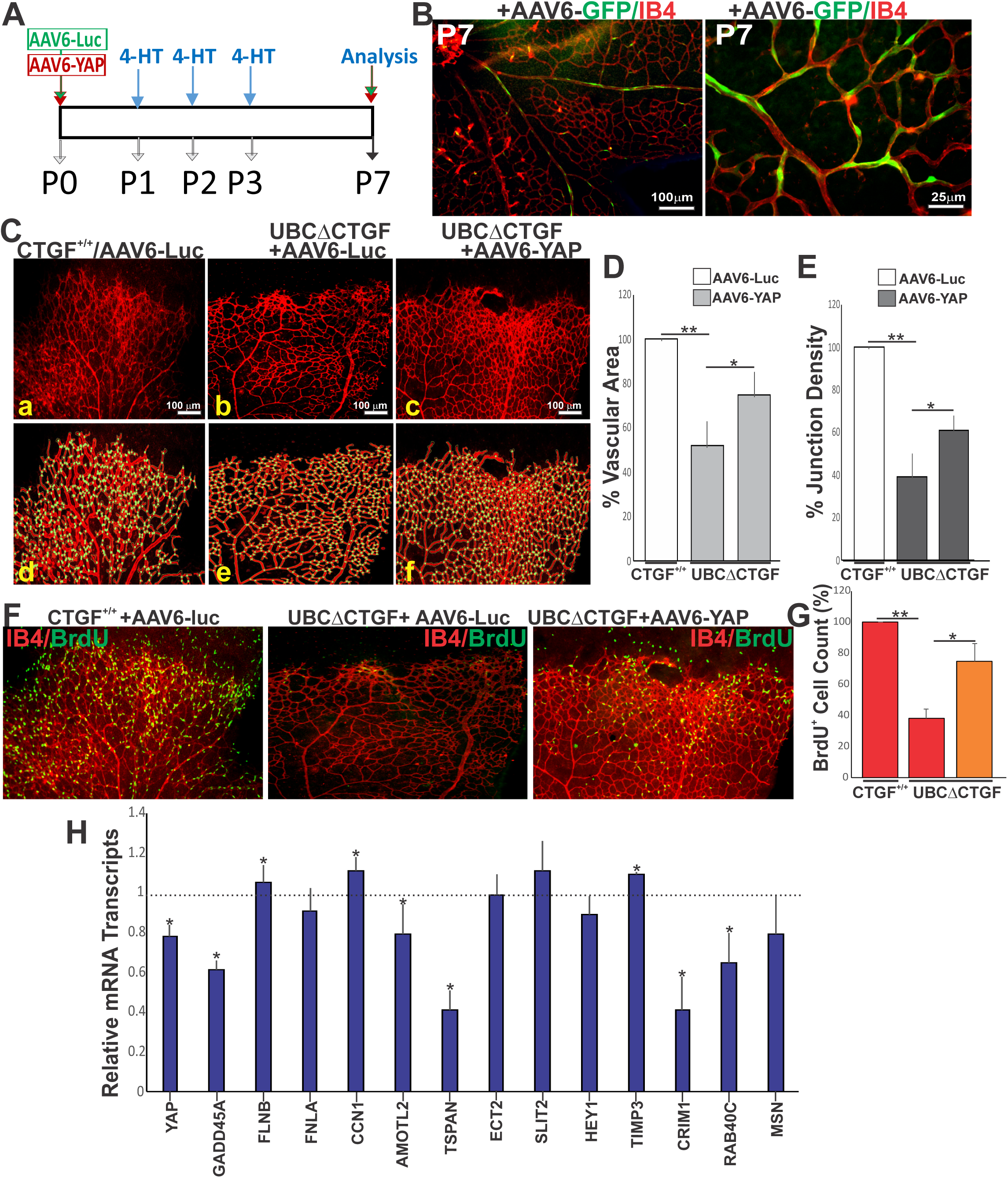
YAP-mediated rescue of vascular defect caused by CTGF deletion. (A) Schematic representation of AAV6-mediated rescue protocol of CTGF deletion following AAV6-mediated expression of YAP. The AAV6 vector is injected at P1 followed by 4HT injection at P2, P3, and P4 to induce CTGF deletion. Vascular phenotype is analyzed at P7. (B) Flat-mount images of IB4-stained retinas from CTGF^+/+^ mice at P7 following retro-orbital injection of AAV6-GFP. Note that GFP expression is largely found in the retinal vascular endothelium lining blood vessels. (C) IB4-stained retinal mounts of CTGF^+/+^ (a) and UBCΔCTGF (b, c) mice following retro-orbital injection of either AAV6-luc or AAV6-YAP. (D-E) Quantitative analyses of vascular parameters performed with AngioTool software as described in Fig. 2 (d-f). (D-E) Graphical representations of the changes in vascular surface and junction density of control and experimental groups described in (B). **, p <0.001 and *, p <0.05 *vs* CTGF^+/+^ (n=4). (F-G) Retinal mount images of BrdU incorporation (*green*) and IB4 staining (*red*) at P7 of CTGF^+/+^ and UBCΔCTGF injected with either AAV6-luc or AAV6-YAP. Proliferation index as determined by counting BrdU^+^ nuclei is shown in (G). Equivalent areas of retinas of control and mutant mice were analyzed. Data are means ± S.E. *, *p <*0.05. (H) Relative mRNA levels of YAP target genes in retinal lysates from CTGF^+/+^ and UBCΔCTGF mouse retinas. The mRNA levels in CTGF^+/+^ mice were set to 1. Each measurement was performed in triplicate. (*, p <0.05, *n*=3).

### Regulation of the Paracellular Vascular Barrier through CTGF-YAP Functional Interaction

Next, we determined whether YAP re-expression may also improve vascular permeability defects associated with CTGF deletion. As shown in Fig. 7A, FITC-albumin extravasation was markedly reduced in UBCΔCTGF mouse retina transduced with AAV6-YAP compared to those transduced with the AAV control. When normalized to the vascular area, re-expression of YAP significantly reduced vascular leakage in CTGF mutant mouse retinas compared to their control counterparts (Fig. 7B).

**Fig. 7.**
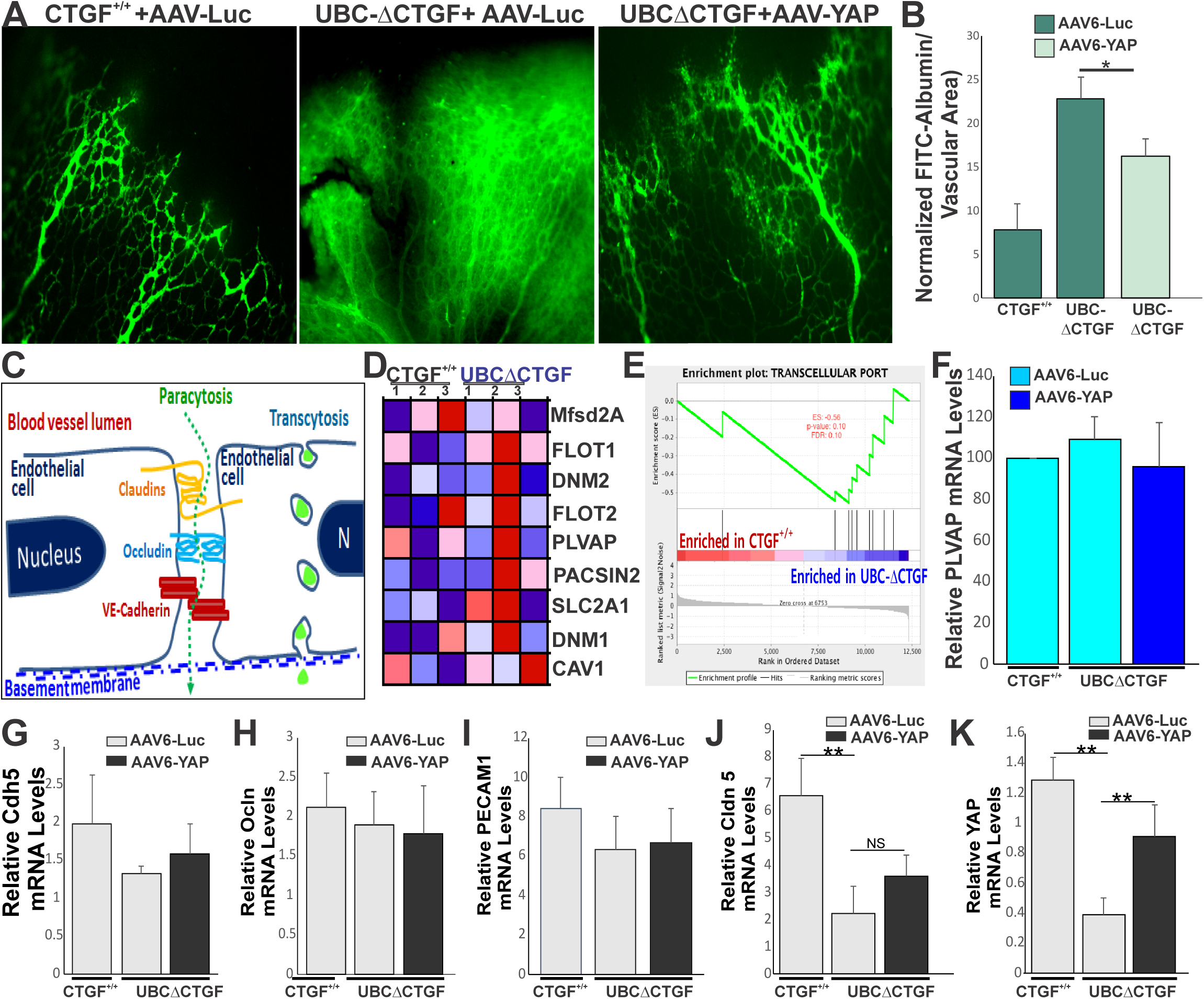
Alteration of paracellular barrier in CTGF-deficient retina. (A) Effects of AAV6-mediated expression of YAP on retinal vascular barrier properties of CTGF^+/+^ and UBCΔCTGF mice. AAV vector injection and 4HT-induced CTGF exon recombination were as described in Fig. 6A. Extravasation of FITC-albumin in retinal flat mounts of CTGF^+/+^ and UBCΔCTGF is shown in (A) and vascular permeability index is shown in (B). (C) Diagram outlining paracellular and transcellular transports in the endothelium. (D-E) GSEA-generated heatmap of CTGF transcriptome enrichment with transcellular transport proteins. Range of colors (red to blue) indicates the range of expression values (high to low). The enrichment score (ES) plot for transcellular gene set is shown in (E). (F-K) Relative mRNA levels of transcellular (e.g., PLVAP) and paracellular (e.g., Cdh5, OCLN, PECAM, CLDN5 and YAP) in retinal lysates from CTGF^+/+^ and UBCΔCTGF mouse retinas. Each experimental point was performed in triplicate. (**, p <0.05, *n*=3).

A defining feature of a functional retinal barrier is (i) a low rate of receptor- and transporter-mediated endocytosis (i.e., transcellular permeability) across ECs, and (ii) the expression of tight and adherens junction proteins that limit paracellular permeability between adjacent ECs (Fig. 7C). Proteins involved in transcellular (endocytic) transport include plasmalemma vesicle-associated protein (PLVAP or PV-1), dynamin-1 and -2, pacsin 2, flotillin-1 and -2, Msfd2a, and the glucose transporter Glut1 (Blumling Iii and Silva, 2012; Bosma et al., 2018). Our GSEA data showed that the CTGF transcriptome is not enriched with these transcellular permeability genes (Fig. 7D-E). Quantitative real time PCR for PLVAP, one of the most important endothelial genes encoding a structural protein of fenestral and stomatal diaphragms (van der Wijk et al., 2019), showed that the low PLVAP levels in WT mouse retinas were not affected by either loss of CTGF function or YAP re-expression (Fig. 7F) suggesting that CTGF signals do not target transcellular transport proteins.

Next, we examined the effects of CTGF signals on EC-EC junctional proteins that regulate the paracellular barrier in the retina. Quantitative analyses of various cell-cell junction proteins showed that the expression of cadherin 5, ZO-1 and occludin was not altered in UBCΔCTGF compared to CTGF^+/+^ retinas, which is consistent with the RNA-Seq data (Fig. 7G-J). Re-expression of YAP in UBCΔCTGF mice did not influence the transcription of these genes, either. CTGF deletion significantly reduced Claudin-5 expression only although ectopic YAP expression had no effect on Claudin 5 transcription in the retina. Claudin-5 deficiency in mice was reported to regulate vascular barrier to small (<0.8 kDa) molecules only (Nitta et al., 2003). As such, changes in Claudin-5 levels do not account for the vascular barrier breakdown in CTGF-deficient mice. Importantly, YAP is also a component of adherens junction complexes and the remarkable reduction of YAP expression in CTGF-deficient mice is potentially responsible for the associated vascular leakage (Fig. 7K).

### EC-EC Interactions Are Stabilized by YAP in CTGF-Deficient ECs

YAP has been shown to associate with the junctional protein complexes in stabilized TJs and AJs (Giampietro et al., 2015; Neto et al., 2018). To examine the role of the CTGF/YAP axis in the regulation of the paracellular barrier, we derived ECs from the brain tissue of UBCΔCTGF mice and their wild-type littermates and cultured them to confluence for subsequent *in vitro* experiments. Under normal culture conditions, CTGF-deficient ECs showed reduced levels of YAP at the mRNA and protein levels (Fig. 8A-C). BrdU-positive cell count was significantly reduced in CTGF-deficient ECs, which is consistent with the role of CTGF and YAP in promoting EC proliferation (Fig. 8B). In CTGF^+/+^-derived cells, YAP localized to the cytoplasm, nucleus and cell-cell junctions. Co-immunoprecipitation and Western blot analyses revealed that YAP forms a complex containing VE-cadherin and α-catenin, corroborating the association of YAP with junctional protein complexes. The levels of YAP-containing protein complexes were reduced in UBCΔCTGF-derived cells. Furthermore, phalloidin-labeled, ZO1-stained cells showed that CTGF mutant ECs exhibit numerous intercellular gaps wherein cell borders (delineated by ZO1 staining) change from straight to zigzag lines (Fig. 8D), which is indicative of and consistent with disrupted cell-cell junctions. Adenovirus-mediated re-expression of YAP, at least in part, reduced intercellular gaps and re-established continuous cell-cell demarcation lines. Together, these observations suggest that CTGF-induced YAP expression is, at least in part, critical for both EC growth and formation and maintenance of stable junctional complexes at cell-cell contacts.

**Fig. 8.**
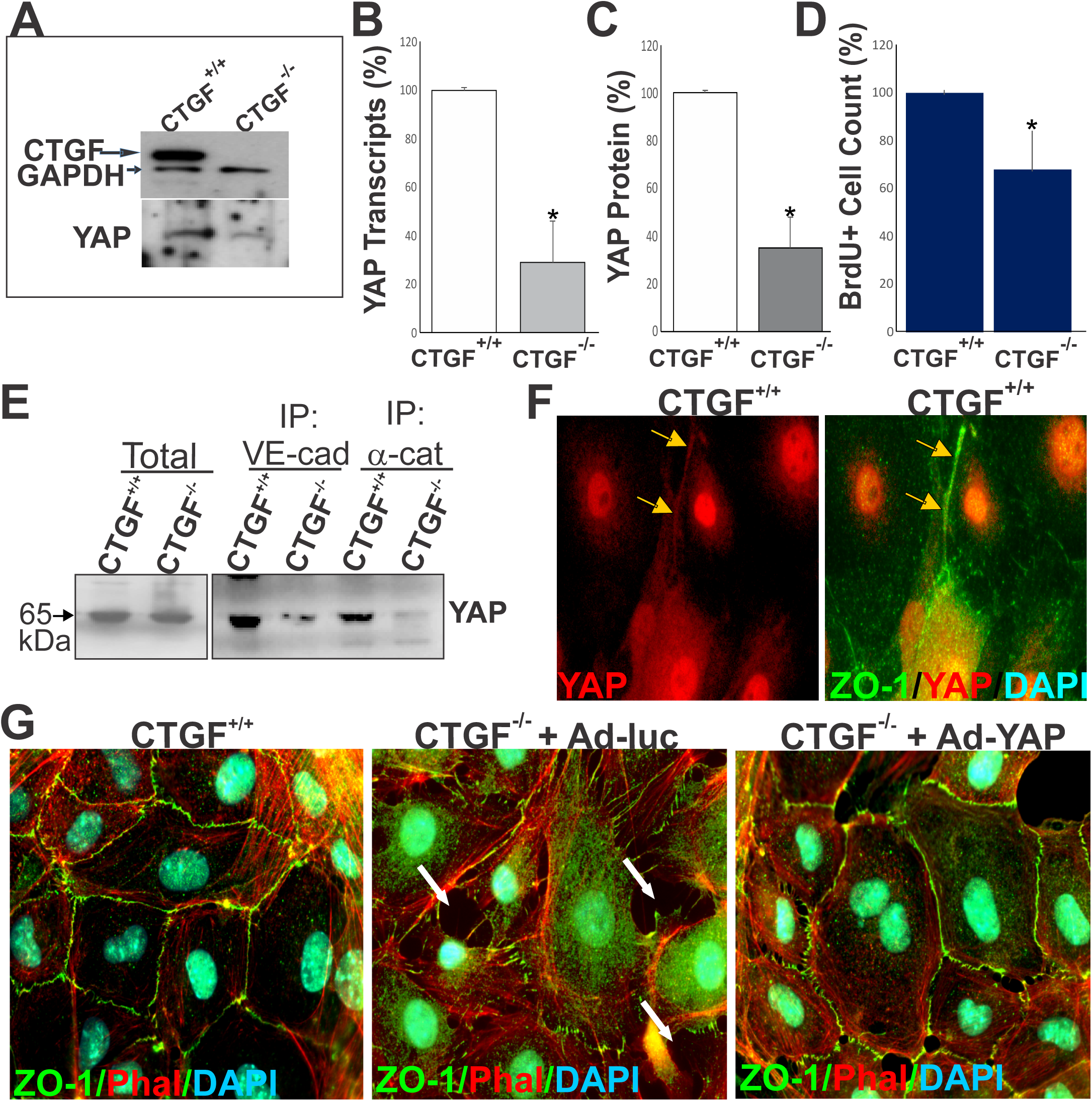
EC-EC interactions are stabilized by YAP in CTGF-deficient ECs. (A) Expression of CTGF and YAP proteins in ECs explanted from CTGF^+/+^ and UBCΔCTGF (i.e., CTGF^-/-^) mouse brain as determined by Western blotting. GAPDH was used as a loading control. (B-C) Relative mRNA and protein levels of YAP in cell lysates from CTGF^+/+^ and CTGF^-/-^ mice. *, p<0.05, n=3. (D) Proliferation index of ECs cultured from CTGF^+/+^ and CTGF^-/-^ mice as determined by cell count. *, p<0.05; n=3. (E) Immunohistochemical localization of YAP in ECs explanted from CTGF^+/+^ mice. Colocalization with ZO-1 is shown. (F) Immunoprecipitation of protein extracts with anti-Cdh5 and anti-α−catenin, and further detection with antibodies against YAP by western immunoblotting. (G) immunohistochemical staining of ECs explanted from CTGF^+/+^ and CTGF^-/-^ mice with phaloidin and ZO-1 antibody. Cells were grown to confluence before immunostaining. Arrows indicate location of intercellular gaps at cell-cell junctions. A representative image of adenovirus-mediated re-expression of YAP is shown.

## DISCUSSION

The matricellular protein CTGF is known to achieve numerous cell type-specific functions in the immediate microenvironment of the cells. CTGF signals regulate cell proliferation, differentiation, adhesion, migration and epithelial mesenchymal transition (Krupska et al., 2015; Leask, 2019). These effects are mediated by a CTGF interactome that includes integrin and non-integrin receptors, other cell-surface molecules (e.g., HSPG, LRPs), ECM and growth factors (Lau, 2016). CTGF-integrin interactions induce angiogenesis in rat cornea (Babic et al., 1999) and in chick chorioallantoic membranes (Shimo et al., 1999), but CTGF was also shown to inhibit angiogenesis *in vitro* by physically interacting with and sequestering VEGF in an inactive form (Inoki et al., 2002). However, while these *in vitro* and ex vivo studies have provided insightful information about the molecular properties of the CTGF protein, they do not integrate exogenous cues such as exposure to heterogeneous cell-cell interactions, flow and associated shear stress effects, as well as tissue-derived signals.

Using transgenic mouse models, targeted gene disruption, and “omic” approaches, we showed that CTGF provides a unique microenvironment in the vascular wall that fine tunes angiogenesis and ensures barrier function. We showed that CTGF is robustly expressed primarily in endothelial stalk cells and secondarily in pericytes and glia during postnatal retinal vascular development and its expression persists in the adult vasculature. In ECs, the expression is limited to cells making new cell-cell contacts while ECs prone to occupy the tip cell position were CTGF-negative. Similarly, retinal neuronal cells such as photoreceptors and ganglion cells minimally, if at all, produce CTGF. Clearly, cells that produce VEGF (e.g., tip cells and astrocytes) do not produce CTGF even though VEGF is known as a major stimulus of CTGF gene expression. The CTGF gene is responsive to a wide range of extracellular stimuli including growth, inflammatory, and stress factors. Hemodynamic forces, which play a critical role in determining the synthetic phenotype and function of ECs, are major inducers of CTGF gene expression (Chaqour et al., 2006; Lee et al., 2018). Tip cells at the sprouting vascular front experience very low levels of shear (Bernabeu et al., 2014) and produce little or no CTGF. Thus, CTGF production could be a direct effect of shear deformation imparted by blood flow in newly lumenized vessels to increase EC capacity to orient in the direction of blood flow. CTGF is indeed involved in the immediate-early response/mechanisms of mechanotransduction in response to flow patterns with or without a clear direction (Chaqour and Goppelt-Struebe, 2006; Hinkel et al., 2014). However, the role of locally produced or blood-borne growth factors in the induction of CTGF gene expression during development cannot be ruled out. Future studies will help elucidate the *in vivo* regulation of CTGF gene expression.

Our data demonstrate that global or EC-specific deletion of CTGF altered normal retinal vascular development and vascular homeostasis of the adult vasculature. Conversely, loss of pericyte-derived CTGF minimally altered vascular growth and integrity indicating the importance CTGF autocrine effects on ECs. Loss of CTGF function at postnatal stages caused hypovascularization of the retina and altered vascular permeability, recapitulating the vascular defects characteristic of early-onset familial exudative vitreoretinopathy in humans (Kramer et al., 2016). CTGF deficiency reduced vascular area and density and junctional point numbers, and increased vascular lacunarity, which correlated with a significant decrease of EC proliferation. However, CTGF deficiency did not alter tip cell number and hierarchical specification of the vasculature into arteries, capillaries and veins, indicating that CTGF expression does not affect the dynamics of EC phenotypical plasticity and specification. Interestingly, global or EC-specific deletion of CTGF produced vascular defects similar to those of transcription factors such as SRF and MRTF-A (Weinl et al., 2015), two versatile transcription factors that toggle between disparate programs of gene expression related to cell growth and differentiation (Sun et al., 2006). The CTGF gene is one of the CArG-box-containing promoter genes (i.e., CarGome) that is activated by the SRF/MRTF duo. Thus, CTGF potentially act as an SRF/MRTF downstream effector molecule mediating their regulatory activity of EC growth and function. Interestingly, the vascular defects of CTGF-mutant mice are distinct from those caused by disruptions of other ECM proteins structurally related to CTGF. For example, deficiency for CCN1, which shares 56% amino acid homology with the CTGF protein, caused retinal vessels to coalesce into large flat hyperplastic sinuses with subsequent loss of their hierarchical organization into arteries, capillaries, and veins (Chintala et al., 2015). CCN1-EC crosstalk controls, directly via integrin signaling and indirectly by fine tuning VEGF signaling, the phenotypical plasticity of ECs and vessel morphogenesis. A previous study by Hall-Glenn et al showed that global CTGF deficiency in mice induced minor enlargement of blood vessels and local edema in the CTGF mutant dermis (Hall-Glenn et al., 2012). This was attributed to incomplete coverage of microvessels by pericytes in the dermis of CTGF mutant mice. However, it was unclear whether the observed vascular phenotype was due to direct effects in ECs or secondary to CTGF effects in other cell types. In our study we show for the first time that the angiogenic defects associated with CTGF deletion were largely due to loss of EC-derived CTGF as EC-specific deletion of CTGF recapitulated, at least in part, those of ubiquitous deletion of CTGF. Because loss of CTGF function reduced EC proliferation, and sprouting, it appears to potentiate the effects of other regulators of angiogenesis such as VEGF.

Molecularly, several clusters of genes and pathways involved in the regulation of important processes such as EC proliferation, migration, and junction formation were affected by CTGF loss of function. The expression of IGF-1, TGF-β1, TGF-β2, FGF-10, and angiopoietin 1 was downregulated upon CTGF deletion and reduced expression of these growth factors may account for the subsequent hypovascularization of the retina. Similarly, loss of CTGF function decreased the expression of several key ECM proteins such as laminin 4, SPARC, and tenascin C as well as other FACIT collagens and proteoglycans, many of which are important for the structural and compliance properties of the vasculature. Thus, CTGF signals potentially regulate the mechanical properties of the vasculature (Chaqour, 2019). Further studies of vascular stiffness regulation by CTGF are warranted.

Additionally, several transcription factors and co-cofactors showed differential gene expression patterns between WT and mutant CTGF mouse retinas including YAP, TEAD, NFAT5, and SRF. CTGF deletion significantly decreased the expression of these transcriptional regulators, many of which affect important aspects of blood vessel stabilization, maturation, and remoledling. YAP regulation by CTGF signals is of a particular significance since, thus far, YAP has been shown to be regulated largely at the post-translational levels by G protein coupled receptors, mechanical forces, and growth factors such as VEGF, via Src family kinases, Rho GTPase, actin dynamics, and Lats1 activity (Aragona et al., 2013; Avruch et al., 2012). CTGF regulation of YAP gene expression adds another level of control to YAP activity.

The regulation of YAP expression by CTGF is a major outcome of our study design although the underlying transcriptional or posttranscriptional mechanisms involved remain to be determined. As a target of YAP and upstream regulator of YAP expression, CTGF appears to reinforce a positive feed-forward loop sustaining vascular development and remodeling. The autonomous role of YAP on vessel development has previously been studied in the mouse retina with EC-specific deletion of YAP and/or TAZ, a paralogue of YAP that also regulates gene expression through binding to YAP interacting partners (Hong and Guan, 2012). The compounded deletion of both YAP and TAZ in the endothelium decreased vascular density, branching and sprouting (Kim et al., 2017), mimicking, at least in part, the CTGF loss of function phenotype. At the functional level, comparing the transcriptomic data of CTGF and YAP signature genes by GSEA showed a significant gene profile overlap of CTGF and YAP targets. YAP plays an important role in cell decisions of proliferation, survival, apoptosis, and adaptive responses to physical and chemical injuries (Zhu et al., 2015). YAP functions to reinforce the cell cytoskeleton and contractile apparatus, and through these mechanisms, YAP appears to control actomyosin-generated internal forces in the cells, thus substantiating the importance of YAP in the geometric control of cell survival and proliferation (Neto et al., 2018). YAP physically interacts with DNA binding transcription factors such as RUNX family, thyroid TF1 (TTF-1), TBX5, PAX3, PAX8, peroxisome proliferator-activated receptor γ, and TEAD to stimulate downstream target gene expression (Panciera et al., 2017). The TEAD consensus sequences were found in at least 75% of YAP target genes. Accordingly, TEAD is required for YAP-induced cell growth, and epithelial–mesenchymal transition. In fact, YAP and TEAD1 co-occupy >80% of the promoters pulled down by chromatin-immunoprecipitation (Zanconato et al., 2015). However, TEAD binding-defective YAP-S94A mutant can still induce expression of a fraction of the YAP-regulated genes (Zhao et al., 2008) indicating that the activities of YAP binding partners is not dependent exclusively on YAP. It is not surprising that while YAP gene expression was downregulated upon CTGF deletion, the expression of TEAD was upregulated in CTGF-deficient mouse retina. Interestingly, CTGF has been identified as a direct TEAD-YAP target gene important for cancer cell growth (Wang et al., 2016). Thus, a self-sustained feedforward loop between CTGF and YAP appears to be involved in vascular development.

Another important finding of our study design is that CTGF is an important determinant of blood vessel integrity and stability. Barriergenesis in the developing retina involves the coordinated induction of proteins that maintain tight and adherens junctions and nutrient transporters in ECs for sprouting blood vessels (Obermeier et al., 2013). Thereafter, maturation of the barrier properties is achieved as functional and physical interactions occur with pericytes, astrocytes, and Muller cells. Assembly and maturation of junctional complexes between ECs are dependent on actomyosin dynamics regulated by the activity of small Rho GTPases such as RhoA and Cdc42 (Aoyama et al., 2012) which appears to be required for maintenance of blood barrier integrity during adulthood as well. However, the acquisition of endothelial barrier properties in CTGF-deficient retinas was not due to defects in transcellular transport since mutant CTGF mice showed no differences in the expression of PLVAP and other membrane proteins that localize to fenestrae and caveolar stomatal diaphragms of retinal vessels. Instead, CTGF deficiency disrupted retinal barrier function through alteration of paracellular barrier protein association and organization. CTGF deletion similarly altered the barrier properties of preformed vessels in adult mice suggesting that basal expression of CTGF is critical for barrier formation, regeneration and maintenance. CTGF effects on the vascular permeability were, at least in part, dependent on YAP expression. Based on CTGF loss- and subsequent YAP gain-of-function phenotypes, reduced YAP levels in ECs disrupted intercellular junctional complex formation and stabilization. Our coimmunoprecipitation experiments showed the presence of YAP in protein complexes of cadherin and α-catenin in agreement with previous findings that YAP associates with the VE–cadherin complex via 14-3-3 proteins, a family of phosphoserine-binding proteins, that inhibit YAP transcriptional activity by preventing its shuttling between the cytoplasm and nucleus (Giampietro et al., 2015). Hyperphosphorylated YAP localizes to adherent junctions in a trimeric complex with 14–3-3 and α-catenin indicating that YAP functions in the cytoplasm, in the nucleus, and at cell-cell junctions. We suggest that the ‘‘decision’’ of the endothelium to become permeable instead of forming a blood retinal barrier is primarily dependent on YAP levels in different cell compartments. It is anticipated that conditions that perturb the functional interactions between CTGF and YAP likewise alters vascular permeability. The full details of the molecular intermediates involved in cellular distribution of YAP remain to be determined.

Taken together, our data showed a crucial role for CTGF in regulating the expression and function of a wide array of extracellular and intracellular factors that are essential for normal vascularization of the retina and for the acquisition of a functional barrier. It is conceivable that genes and expression clusters affected by CTGF in the endothelium might be relevant for angiogenesis and barriergenesis in organs other than the retina. It also is highly likely that the deregulation of the CTGF network genes may be relevant for processes of pathological angiogenesis, which are characterized by disrupted growth, lack of vessel maturation, defective remodeling and/or barrier dysfunction. Future studies focused on the characterization of the relevance of this novel CTGF-based angio-modulatory pathway in *in vivo* models of vascular diseases are warranted.

## EXPERIMENTAL PROCEDURES

### KEY RESOURCE TABLE

| REAGENTS or RESOURCE | SOURCE | IDENTIFIER |
| --- | --- | --- |
| Experimental Models : Mice |  |  |
| Tg(Ctgf-EGFP)FX156Gsat/Mmucd | MMRRC | 011899-UCD |
| CTGF<sup>flox/flox</sup> | Liu et al, 2011 | N/A |
| Tg(UBC-cre/ER<sup>T2</sup>)1Ejb | Jackson Labs | 007001 |
| Tg(Cdh5-PAC-cre/ER<sup>T2</sup>) | Sorensen et al, 2009 | N/A |
| Tg(*Cspg4-cre*/ER<sup>TM</sup>) | Jackson Labs | 008538 |
| Experimental Models: Cells |  |  |
| HRECs | Cell Biologics | H6065 |
| Brain/Aorta ECs | This paper | N/A |
| Recombinant viruses |  |  |
| AAV6-GFP | GTRP-U Penn | N/A |
| AAV6-Luc | GTRP-U Penn | N/A |
| AAV6-YAP | Vigene Bioscience | N/A |
| Ad-luc | GTRP-U Penn | N/A |
| Ad-YAP | Creative-Biogene | N/A |
| Antibodies |  |  |
| Collagen IV | Millipore Sigma | Cat# AB756P |
| Iba-1 | Wako | Cat# 019-19741 |
| NG2 | Millipore Sigma | Cat# MAB5384 |
| ZO-1 | Invitrogen | Cat# 61-7300 |
| Phospho YAP (Ser127) | Cell signaling | Cat# 4911 |
| YAP | Santa Cruz | Cat# sc-101199 |
| VE-Cadherin | Santa Cruz | Cat# sc-9989 |
| a-catenin | Abcam | Cat# Ab51032 |
| β-Catenin | Cell Signaling | Cat# 9562 |
| GFAP | Biovision | Cat# A1124-50 |
| F4/80 | BIO-RAD | Cat# MCA497 |
| Conjugated mouse IgG | Cell signaling | Cat# 7076S |
| *Conjugated rabbit IgG* | Cell signaling | Cat# 7074S |
| Deposited data |  |  |
| Raw and analyzed data | This paper | GEO: GSE146900 |
| Chemicals |  |  |
| Isolectin B4 | Vector Laboratories | Cat# DL-1207 |
| DAPI | Millipore Sigma | Cat# F6057 |
| FITC-albumin | Millipore Sigma | Cat# A9771 |
| Phalloidin | Cytoskeleton | Cat# PHDR1 |
| Software and Algorithms |  |  |
| Angiotool | Zudaire et al, 2011 | N/A |
| Hisat2 software | Kim et al, 2015a | N/A |
| StringTie | Pertea et al, 2015 | N/A |
| Ballgown R package | Frazee et al., 2015 | N/A |

#### Mice

GENSAT Tg(Ctgf-EGFP)FX156Gsat/Mmucd (011899-UCD) mice, referred to herein as CTGF-GFP mice carrying enhanced green fluorescent protein (GFP) under the control of the CTGF promoter, were developed under the NINDS-funded GENSAT BAC transgenic project (Gong et al., 2003) and obtained from the Mutant Mouse Regional Resource Center. CTGF-GFP mice were initially in the FVB/N-Swiss Webster background and later backcrossed for >10 times in the C57BL/6J genetic background. CTGF^flox/flox^ were previously described (Liu et al., 2011). Tg(UBC-cre/ER^T2^)1Ejb, Tg(Cdh5-PAC-cre/ER^T2^) and Tg*(Cspg4-cre/ER™*) mice were from Jackson Laboratory (Ruzankina et al., 2007; Sorensen et al., 2009; Zhu et al., 2011). All animal studies were carried out in accordance with the recommendations in the Guide for the Care and Use of Laboratory Animals of the National Institutes of Health. Mice were handled and housed according to the approved Institutional Animal Care and Use Committee (IACUC) protocol 14-10425 of SUNY Downstate Medical Center, NY, USA.

### Generation of the CTGF conditional allele

For global and cell-specific removal of floxed STOP-cassettes, CTGF^flox/flox^ were crossbred with Tg(UBC-cre/ER)1Ejb, Tg(Cdh5-PAC-cre/ER^T2^) and Tg(Cspg4*-*cre/ER™) mice to induce global, EC-, and pericyte-specific deletion of CTGF respectively. Mouse genotypes were determined by PCR to identify mice with floxed alleles and hemi-and homozygous floxed alleles with or without the Cre allele. A solution of 4-hydroxytamoxifen (4HT) was dissolved in ethanol at 10 mg/ml, and then 4 volumes of corn oil were added. Samples of 4HT were thawed and diluted in corn oil prior to intraperitoneal injection of 100 μl to mouse pups. The relative recombination levels in mutant mice as compared with CTGF^+/+^ were determined as previously described previously (Liu et al., 2011).

### Recombinant AAV vector generation and retroorbital injection

AAV6 vectors expressing GFP or YAP cDNA under the control of the cytomegalovirus (CMV) promoter were used. AAV6-GFP control was obtained from the Penn Vector Core at the University of Pennsylvania. AAV6-YAP vector was produced by triple transfection of 293T cells with the pAV-FH AAV vector, the Ad helper vector, and a vector encoding the Rep and serotype-specific Cap proteins, followed by iodixanol gradient centrifugation and purification by anion-exchange chromatography. Viral titers were determined as the number of genomic copies (GC) per microliter using quantitative real-time PCR (1×10^12^ GC/ml). AAV packaging, purification, and quality control were performed by Vigene Biosciences. For retroorbital injection, mouse pups were anesthetized using ketamine/xylazine mixture prior to retro-orbital injection of the recombinant viral vectors (4×10^12^ vector genomes) in 15 μl of sterile phosphate buffer solution (PBS).

### Cell culture

Human retinal endothelial cells were purchased from Cell Biologics and authenticated by endothelial marker expression (CD31/DiI-acLDL double-positive) and morphology. Cells were cultured in endothelial basal medium containing manufacturer’s supplements. Isolation of mouse brain and aorta ECs was performed as previously described. Primary brain and aorta ECs were isolated by immunopanning from both P7 male and female wild-type or UBCΔCTGF mice. Cells were cultured in endothelial medium supplemented with 10% fetal bovine serum (FBS) to promote formation of confluent monolayer cultures. Cells were switched to media with low serum (2% FBS) in the absence of growth factors after reaching confluence and prior to starting experimental treatments described in the text. All cell cultures were used between passage 3 to 9.

### Adenoviral vector infection

Mouse YAP cDNAs were isolated by PCR amplification using DNA templates obtained from Addgene (Watertown, MA) and cloned into a shuttle vector at Creative-Biogene (Shirley, NY). The recombinant adenovirus, Ad-YAP, was produced by cotransfecting an adenoviral shuttle vector with a viral backbone in which the recombinant cDNA is driven by the CMV promoter. All adenoviruses were replication deficient and used at 20 multiplicity of infection to infect cultured ECs. Adenoviral vectors were added to ECs cultured in medium containing 2% FBS and incubated for 4 h. Thereafter, the cells were washed five times and cultured in medium with 10% FBS and supplements. The adenovirus encoding the luciferase gene, Ad-luc (obtained from the Gene Therapy Resource Program Vector Core Laboratory at the University of Pennsylvania) was used as a control.

### Quantification of junctional density, length, and lacunarity of retinal vessels

Mouse eyes were collected at the indicated postnatal days and fixed in 4% paraformaldehyde (PFA) for 2 h. Retinas were dissected and laid flat on SuperFrost^R^ Plus-coated slides, permeabilized in 0.1% Triton X-100 at room temperature and stained with Isolectin B4 (IB4), as previously described (Lee et al., 2017a). Fields of view of the retinal vascular networks from control and mutant mice were captured by using the x2 and x40 objective lenses. Using AngioTool (Zudaire et al., 2011), the variation in foreground and background pixel mass densities across an image, vascular parameters such as total vascular surface area, vessel junctional density and “gappiness”/lacunarity of the vascular network were determined. For each parameter, at least four fluorescent images/retina were taken from 4–5 mice. The data are presented as means ±S.E. The statistical significance of differences among mean values was determined by one-way analysis of variance and two-tailed *t* test.

### Antibodies and immunohistochemical staining

A set of eyes were frozen in Tissue-Tek Optimal Cutting temperature compound and ten micrometer-thick cryostat sections were prepared. Retinas were dissected from another set of eyes and laid flat on SuperFrost® Plus coated slides to obtain whole mount preparations. Sections and flat mount preparations were then permeabilized in 0.1% Triton X-100 at room temperature for 20 min and further double stained with IB4 and/or with the following antibodies: Iba-1 (1:500, Wako), Collagen IV (1:500, EMD Millipore), NG2 (1:200, EMD Millipore), ZO-1 (1:200, ThermoFisher Scientific) YAP (Cell 1:200, Signaling) or Phalloidin (3.5:500, Cytoskeleton). Immunodetection was performed with either rhodamine or fluorescein-conjugated anti-mouse or anti-rabbit secondary antibody diluted in blocking solution. Retinal mounts and sections were mounted in DAPI. Images were acquired using a Leica DM5500B fluorescence microscope (Leica). To localize proteins in cultured cells, cells were fixed in PFA for 10 min and permeabilized in 0.1% Triton X-100 at room temperature for 5 min. Cells were then incubated with the indicated primary antibodies overnight at 4 °C and treated with TRITC- or FITC-conjugated secondary antibodies. Images were captured using a Leica DM5000 B fluorescence imaging system.

### BrdU incorporation and cell proliferation assay

CTGF mutant and wild-type littermate control animals was intraperitoneally injected with 100 μl of 3 μg/ml BrdU per 3 g body weight. For BrdU labeling, retinas were digested with Proteinase K (10 μg/ml), fixed in 4% PFA, treated with DNase I (0.1 units/ml) for 2 h at 37 °C, and incubated with anti-BrdU antibody (1: 50, BD Pharmingen). ECs were visualized by staining with IB4 (Molecular Probes), and BrdU was detected using conjugated mouse anti-BrdU Alexa 488 (Molecular Probes). BrdU^+^ cells that co-localized with retina vessels were counted, and numbers of BrdU^+^ cells were normalized to the vessel area or vessel length determined by AngioTool. The sum of BrdU^+^ cells for the four leaflets of the retina was normalized to the sum of analyzed vessel area for each animal. For cultured cells, proliferation rate was determined using the CyQUANT Direct Cell Proliferation Assay according to the manufacturer’s protocol (Invitrogen). The fluorescence intensity was measured with a fluorescence microplate reader using an FITC filter set.

### In vivo permeability assays

Vascular permeability is determined using the highly sensitive FITC-albumin permeability assay. Anesthetized mice received retro-orbital injections of 50 μl of 1% albumin-AlexaFluor 488 (Sigma-Aldrich). After 5 min, animals were sacrificed and the eyes were enucleated and fixed in PFA for 10 min. Retinas were dissected from the eyewall and optic nerve and FITC-albumin was immediately visualized with a Leica 65700 fluorescence microscope and analyzed using DFM software. Retinas were further stained with IB4. The area of albumin present in the retina was normalized to that of total vascular areas. For quantitative analysis of the vascular barrier leakage, anesthetized mice received intraperitoneal injections of 100 μl of EB (45 mg/kg). After perfusion with PBS, harvested retinas were harvested, homogenized and incubated in formamide for 24 hours at 55°C. Supernatants were collected and EB was measured at the absorbance of 620 nm. The concentration of dye in the extracts was calculated from a standard curve.

### RNA isolation and RT-qPCR

Total RNA was extracted from mouse retina using TRIzol reagent (Sigma). cDNA synthesis was carried out by using the Prime script reverse transcript kit (Takara). Highly specific primers were designed using Web-based primer design programs. A qPCR was performed with biological triplicates using Power SYBR™ green PCR premix (Applied biosystems). The cycling parameters for qPCR amplification reactions were: AmpliTaq activation at 95°C for 10min, denaturation at 95°C for 15s, and annealing/extension at 60°C for 1min (40 cycles). Triplicate *Ct* values were analyzed with Microsoft Excel using the comparative *Ct* (ΔΔ^Ct^) method as described by the manufacturer. The transcript amount (determined by the − 2^ΔΔCt^ threshold cycle [C_T_] method) was obtained by normalizing to an endogenous reference (18S rRNA) relative to a calibrator.

### Library preparation for Illumina sequencing

Retinas were harvested for CTGF^+/+^ and UBCΔCTGF at P7 (3 replicates from each control and mutant mice originating from different litters) and total RNA extracted using TRizol. 1∼2 μg total RNA of each sample was taken for RNA-seq library preparation. RNA-Seq analysis was performed by Arraystar (USA). Briefly, mRNA was isolated from total RNA with NEBNext® Poly(A) mRNA Magnetic Isolation Module. The enriched mRNA was used for RNA-Seq library preparation using KAPA Stranded RNA-Seq Library Prep Kit (Illumina). The library preparation procedure included (a), fragmentation of the RNA molecules; (b) reverse transcription to synthesize first strand cDNA; (c) second strand cDNA synthesis incorporating dUTP; (d) end-repair and A-tailing of the double stranded cDNA; (e) Illumina compatible adapter ligation; and (f) PCR amplification and purification for the final RNA-Seq library. The completed libraries were qualified on an Agilent 2100 Bioanalyzer for concentration, fragment size distribution between 400 ∼ 600 bp, and adapter dimer contamination. The amount was determined by the absolute quantification qPCR method. The bar-coded libraries were mixed in equal amounts and used for sequencing. The DNA fragments in well mixed libraries were denatured with 0.1M NaOH to generate single-stranded DNA molecules, loaded onto channels of the flow cell at 8 pM concentration, and amplified *in situ* using TruSeq SR Cluster Kit v3-cBot-HS (#GD-401-3001, Illumina). Sequencing was carried out using the Illumina HiSeq 4000 according to the manufacturer’s instructions. Sequencing was carried out by running 150 cycles.

### Filtering and merging of reads

Image analysis and base calling were performed using Solexa pipeline v1.8 (Off-Line Base Caller software, v1.8). Sequence quality was examined using the FastQC software. The trimmed reads (trimmed 5’,3’-adaptor bases using Cutadapt) were aligned to reference genome using Hisat2 software (v2.0.4) (Kim et al., 2015a). The transcript abundance for each sample was estimated with StringTie (v1.2.3) (Pertea et al., 2015), and the Fragments Per Kilobase of transcript per Million mapped (FPKM) value (Mortazavi et al., 2008) for gene and transcript levels were calculated with R package Ballgown (v2.6.0) (Frazee et al., 2015). The thresholds used for identifying differentially expressed genes (DEGS) were: (i) DESeq. 2 mean normalized counts >10; (ii) padj-value <0.5, and (iii) log_2_fold change >0 (Love et al., 2014). rMATS (Shen et al., 2014) was used to analyze alternative splicing events.

### Pathways analysis

The differentially expressed genes/transcripts were analyzed for their enrichment in gene ontological functions or pathways using the TopGo R software package. The statistical significance of enrichment was given as p-value by Fisher exact test and -log_10_(p) transformed to Enrichment score. Principal Component Analysis (PCA) was performed with genes that have the ANOVA p value < 0.05 on FPKM abundance estimations. Statistical or graphics computing, correlation analysis, Hierarchical Clustering, scatter plots and volcano plots were performed in R, Python or shell environment. The raw RNA-Seq data and analyzed FPKM values can be accessed through NCBI/GEO (GSE146900).

### Gene set enrichment analysis

RNA-Seq dataset expression of gene profiling files downloaded from the dataset was analyzed by Gene Set Enrichment Analysis (GSEA, http://www.broad.mit.edu/gsea/index.html). Gene sets are available from Molecular Signatures DataBase (MolSigDB, http://www.broad.mit.edu/gsea/.msigdb/msigdb_index.html). The whole genome (27455 genes) with expression values were uploaded to the software and compared with catalog C5 gene ontology gene sets in MsigDB (Subramanian et al., 2005), which contains 233 GO cellular component gene sets, 825 GO biological process gene sets, and 396 GO molecular function gene sets. GSEA was run according to default parameters: collapses each probe set into a single gene vector (identified by its HUGO gene symbol), permutation number = 1000, and permutation type = “gene-sets”. Calculation of the false discovery rate (FDR) was used to correct for multiple comparisons and gene set sizes. Heat maps were generated using GENE-E, from the Broad Institute (http://www.broadinstitute.org/cancer/software/GENE-E/) for the CTGF and YAP transcriptomes.

### Western immunoblotting

For protein analysis from retinas, mouse eyes were enucleated and retinas were carefully dissected and homogenized in lysis buffer containing 10 mM NaF, 300 mM NaCl, 50 mM Tris, pH 7.4, 1% Triton X-100, 10% glycerol, and 1mM EDTA with a 1% volume of phosphatase and protease inhibitor mixture. Protein samples (20 μg) were fractioned in a 10% SDS-polyacrylamide gel, transferred to a nitrocellulose membrane, and Western blot analysis was performed with each of the indicated primary antibodies. Immunodetection was performed using enhanced chemiluminescence (ECL) from Pierce. Protein bands were quantified by densitometric scanning. To analyze proteins from cell cultures, cells were homogenized in lysis buffer fractioned by electrophoresis and analyzed as described above. The primary antibodies used in this study were as follows: pYAP (1:500; Cell Signaling), YAP (1:500; Santa Cruz), GAPDH (1:1,000; Santa Cruz). CTGF antibodies (1:500) were generated using two 16-amino acid peptides (residues between 179 and 195 in the primary sequence of the mouse CTGF, GenBank accession no. NM_010217) was synthesized and purified by high-performance liquid chromatography (Biosynthesis). Antibodies were raised against KLH-coupled peptide in rabbits by Pocono Rabbit Farm and Laboratory Inc (Poconos, PA). After completion of the immunization process, the rabbits were exsanguinated, and the antibodies were purified by affinity chromatography using affinity columns. Bound antibodies were eluted with 100 mM glycine and immediately neutralized with Tris base and adjusted to a concentration of 0.4 mg/ml before storage at -20°C. Serum titer was determined by enzyme-linked immunosorbent assay.

### Immunoprecipitation

Total proteins (∼10 mg) of cultured cell lysates were transferred to tubes with antibody-bound protein G beads and rocked gently at 4°C overnight. Non-specific bound proteins were removed with five washes with 1× PBS containing 1% NP-40. Immunoprecipitation products were extracted from the protein G beads using sample loading buffer and were further analyzed by Western immunoblotting.

### Statistical Analyses

Data were expressed as mean ± S.E. To test differences among several means for significance, a one way analysis of variance with the Newman-Keuls multiple comparison test was used. Where appropriate, a *post hoc* unpaired *t* test was used to compare two means/groups. Statistical significance was set to *P* value less than 0.05

## SUPPLEMENTAL INFORMATION

Supplemental Information includes four figures.

## DATA AVAILABILITY

RNA-Seq data are available at the NCBI Gene Expression Omnibus with the series record GSE146900.

## AUTHOR CONTRIBUTIONS

SM and SL generated and characterized the vascular phenotype of mice with tissue-specific deletion of CTGF. JAC and SP performed mouse genotyping and *in vitro* studies, including cell infection and transfection and data analyses. AL generated mice with floxed CTGF exons. JS performed RNA-Seq data analyses and GSEA. JAK contributed to transcriptomic data interpretation. BC designed the project and experimental approaches and wrote the paper.

## ACKNOWLEDGMENTS

We are thankful to Dr Stephen J. Weiss (University of Michigan) for his involvement in the project conception, data discussion and interpretation and review of the manuscript. We appreciate the technical contribution of Genesis Lopez and Xin Chen and the help of Zaid McKie-Krisberg for RNA-Seq data formatting and submission to GEO. We thank all past and present lab members for their contributions to the generation and characterization of genetically modified animals and discussions during the preparation of the manuscript. This work was supported in part by grants from the National Eye Institute of the National Institutes of Health (grants EY024998 and EY022091-05A1) to B.C.

## Notes

https://www.ncbi.nlm.nih.gov/geo/query/acc.cgi?acc=GSE146900

