## Supplemental Figures for "A CTGF-YAP regulatory pathway is essential for angiogenesis and barriergenesis in the retina"

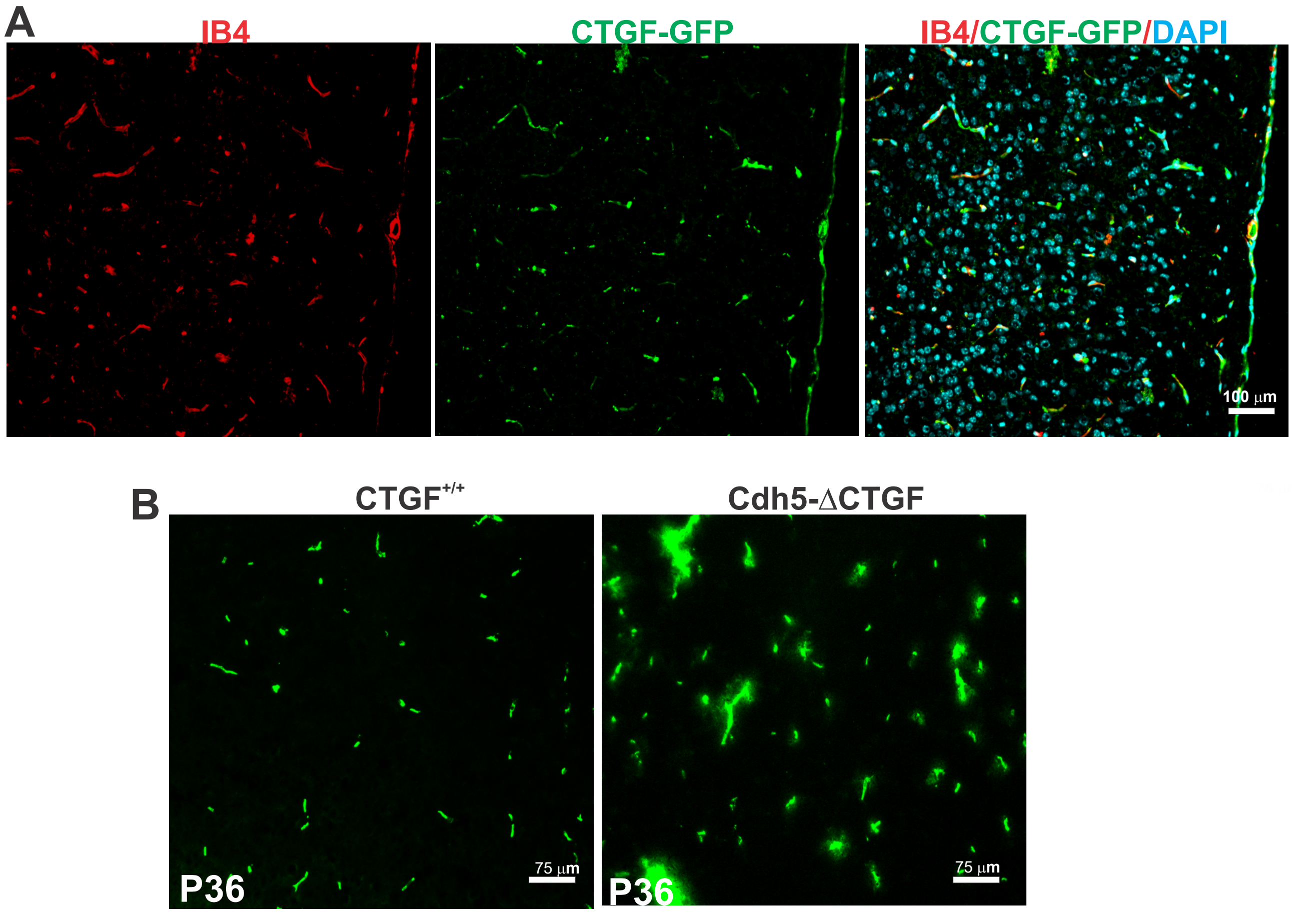


**Fig. S1. CTGF expression and barrier function in mouse brain blood vessels.** (A) Tissue sections of CTGF-GFP mouse brain were labeled with IB4 (red for blood vessels). The merged image indicates that in CNS blood vessels are all positive for GFP (used as a proxy for CTGF expression). (B) Defective blood brain barrier in adult (P36) mice with endothelial deficiency of CTGF. The FITC-albumin tracer was confined to the capillaries in WT littermates, whereas Cdh5ΔCTGF mouse brain showed tracer leakage in the brain parenchyma surrounding capillaries in cortical areas.

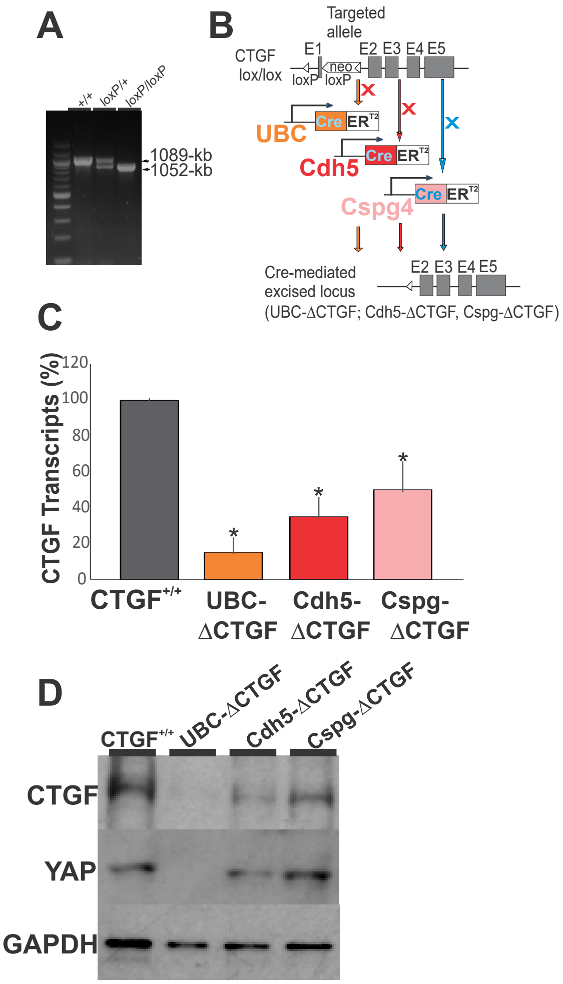


**Fig S2. Targeting of the CTGF genomic locus and recombination in UBC-CreER$^{T2}$, Cdh5(PAC)-CreER$^{T2}$ and Cspg4-CreER$^{T2}$ mice.** (A) CTGF wild-type allele was detected by PCR as a 1052-bp DNA band. The floxed allele was confirmed by the presence of a 1089-bp DNA band. (B) Schematic diagrams showing the targeted CTGF genomic locus with the CTGF flox-neo allele, the CTGF flox allele with the neomycin resistance cassette deleted by recombination performed with loxP sites, and the mutant locus following 4HT-induced UBC-, Cdh5 or Cspg4 promoter-induced Cre-mediated excision. (C) Relative mRNA levels of CTGF in retinal lysates from CTGF$^{+/+}$, UBC-ΔCTGF, Cdh5-ΔCTGF and CspgΔCTGF. Each measurement was performed in triplicate (n=3). *, p <0.01 *vs* CTGF$^{+/+}$ (n=4). (D) CTGF protein detection by Western immunoblotting in retinal lysates from CTGF$^{+/+}$, UBCΔCTGF, Cdh5ΔCTGF and CspgΔCTGF mice following consecutive daily injection of 4HT. GAPDH signal was used as a loading control.

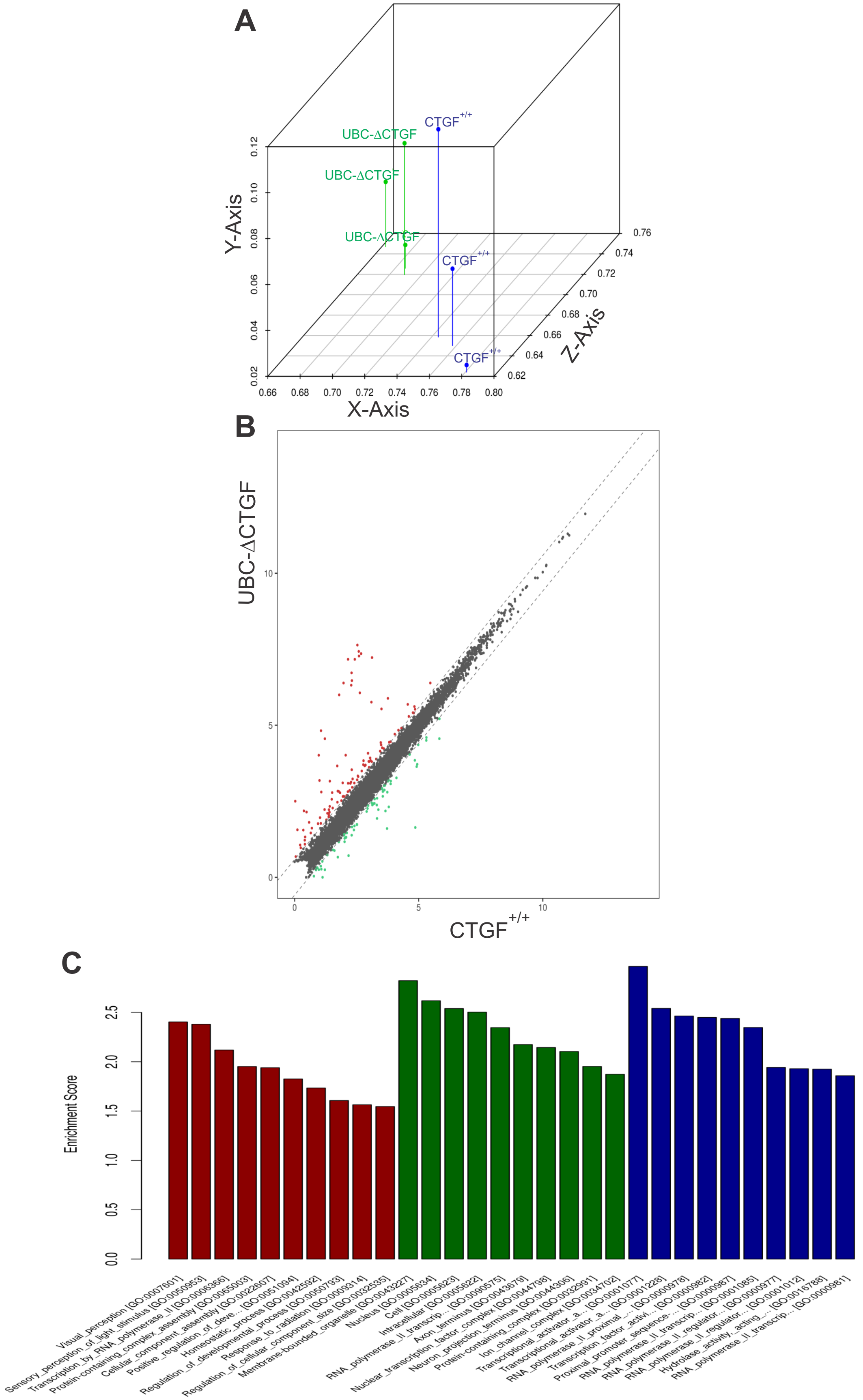


**Fig. S3. Transcriptome profiling of $CTGF^{+/+}$ and UBC-ΔCTGF mouse retina at P7.** (A) PCA analysis showing hallmark gene signatures with clear segregation between $CTGF^{+/+}$ and UBCΔCTGF transcriptomes of the three biological replicates plotted in 3 D volumetric space. X-axis (52.08%), Y-axis (0.41%) and Z-axis (47.51%). (B) Scatter plots comparing cross-sample normalized RNA-Seq read counts of CTGF target genes. Red and green symbols are transcripts with FDR < 0.05 and enrichment >2 fold (values are $log_2$ of normalized transcript counts). (C) Top GO biological process terms enriched in up- or downregulated genes in retinas of CTGF mutant mice at P7.
